# Inhibition of histone acetyltranserase function radiosensitizes CREBBP/EP300 mutants via repression of homologous recombination, potentially targeting a novel gain of function

**DOI:** 10.1101/2020.04.10.028217

**Authors:** Manish Kumar, David Molkentine, Jessica Molkentine, Kathleen Bridges, Tongxin Xie, Liang Yang, Andrew Hefner, Meng Gao, Mitchell J. Frederick, Sahil Seth, Mohamed Abdelhakiem, Beth M. Beadle, Faye Johnson, Jing Wang, Li Shen, Timothy Heffernan, Aakash Sheth, Robert Ferris, Jeffrey N. Myers, Curtis R. Pickering, Heath D. Skinner

## Abstract

Despite radiation forming the curative backbone of over 50% of malignancies, there are no genomically-driven radiation sensitizers for clinical use. We performed *in vivo* shRNA screening to identify targets generally associated with radiation response as well as those exhibiting a genomic dependency. This identified the histone acetyltransferases CREBBP/EP300 as a target for radiosensitization in combination with radiation in cognate mutant tumors. Further *in vitro* and *in vivo* studies confirmed this phenomenon was due to repression of homologous recombination following DNA damage and can be reproduced using chemical inhibition of histone acetyltransferase (HAT), but not bromodomain function. Selected mutations in CREBBP lead to a hyperacetylated state that increases CBP and BRCA1 acetylation, representing a gain of function targets by HAT inhibition. Additionally, mutations in CREBBP/EP300 were associated with recurrence following radiation, in several squamous cell carcinoma cohorts. These findings represent both a novel mechanism of treatment resistance and the potential for genomically-driven treatment.

## Background

With a few isolated exceptions, the cure of solid tumors requires effective local therapy. In the vast majority of disease sites this translates to a need for radiation, either alone or as part of a treatment package including surgery. Despite the large number of novel targeted and immunotherapies introduced over the past decade or more, in the vast majority of cases the only agents available to improve responses to radiation are the cytotoxic chemotherapies that have been in use since the 1980’s or earlier. Because of this, an effective and minimally toxic radiosensitizer has the potential to, in short order, positively impact hundreds of thousands of patients.

One exemplar of this phenomenon is head and neck squamous cell carcinoma (HNSCC), the curative treatment of which has remained largely unchanged over the past two decades. While 75% of patients with HNSCC require radiation for the treatment of their disease, the recent failure of cetuximab means there are generally no biologically-driven radiosensitizers available to improve response and decrease toxicity of this therapy^1–4^. Nor has the advent of immunotherapy changed the current paradigm, with a recent clinical trial of chemoradiation combined with immunotherapy closed due to lack of efficacy^5^.

Despite these failures, the search for improved combinations with radiation remains critical. Again, focusing solely on the close to 300,000 patients with HNSCC annually worldwide who are recommended to receive radiation, an agent that improves the efficacy of this treatment by 15% could lead to more than as 30,000 lives saved annually ^6,7^.

However, most anti-neoplastic agents tested in the pre-clinical setting ultimately fail to be translated to the clinic due to multiple factors, including the artificial nature of *in vitro* systems and unforeseen toxicity^8,9^. Targets identified as radiosensitizers in an *in vitro* model, may underperform *in vivo* due to complex interactions within the tumor itself. Additionally, the same microenvironment interactions could be potential targets for radiosensitization and may not be readily identified using *in vitro* screening techniques.

Additionally, most large-scale screening approaches using cell lines with known genomic status have not exposed the cells to radiation and, thus, have not identified tumor mutations or alterations that may be associated with specific targets for radiosensitization^10^. The model of a genomically-mediated “Achilles heel” in tumors harboring a specific genotype has perhaps best been characterized for BRCA1-altered tumors and their dramatic response to PARP inhibition^11^. However, this model can be expanded to the combination of novel agents with DNA-damaging therapies in specific genetic backgrounds^12^. This is highly advantageous as genomically-driven radiosensitizers have the potential to only affect the mutated cancer cells, and not normal cells, providing improved responses with less toxicity.

In the current investigation, we began with an *in vivo* screening analysis to identify novel radiosensitizers in HNSCC, with a further analysis to identify potential targets for based upon somatic mutation. These data led to the identification of histone acetylation as a novel target, potentially driven by a gain of function in certain classes of HAT/TAZ2 domain mutants. These mutations exhibit reduced basal inhibitory function, leading to a hyperacetylated state and potential dependency on homologous recombination and BRCA1 for DNA damage repair. The importance of CREBBP and

EP300 mutation were underscored following analysis of tumor tissues in several cohorts of patients with SCC of the head and neck, lung or cervix treated with radiation therapy, identifying these mutations as associated with radioresistance and poor outcome.

## Results

### In vivo screening identifies multiple potential radiosensitization targets

We performed *in vivo* shRNA library screens in tumors generated from 5 HNSCC cell lines of varying genetic background and HPV status (HPV-positive: UM-SCC-47 & UPCI-SCC-152), HPV-negative: UM-SCC-22a, HN31 & Cal 27) treated with radiation (Supplementary Table 1). Two libraries were used: one encompassing most genes known to be targeted by available anti-neoplastic agents currently in clinical use or in clinical trials and the second targeting the DNA-damage repair pathway (Supplementary Table 2).

To determine a gene level summary estimate of the impact of knock-out of each gene, we performed redundant shRNA analysis (RSA) to generate log p-values for each gene for irradiated tumors (Fig. 1A). Additionally, quantile transformed median fold change (fc) for each target was analyzed (Fig. 1B). This analysis identified several targets known to be associated with radiosensitization, such as CDK6, XIAP, PIK3CA and PTK2^13-17^. However, to further identify targets specific for radiation response, as opposed to those primarily inhibiting tumorigenesis, we evaluated RSA-log p values for irradiated compared to untreated tumors in complimentary experiments reported elsewhere^18^ (Fig. 1 C-D). We examined the average for each group and selected targets highlighted in red in Fig. 1D, based on values favoring radiation response versus effects on tumorigenesis, defined very broadly to maximize potential target identification (Supplementary Table 3 for selected targets). In this group, we identified several multiple DNA damage repair genes, such as TP53BP1, ATRIP and RUVBL1. Additionally, inhibition of XIAP and PI3KCA were highly associated with radioresponse, even above their anti-tumor effects.

**Fig. 1:**
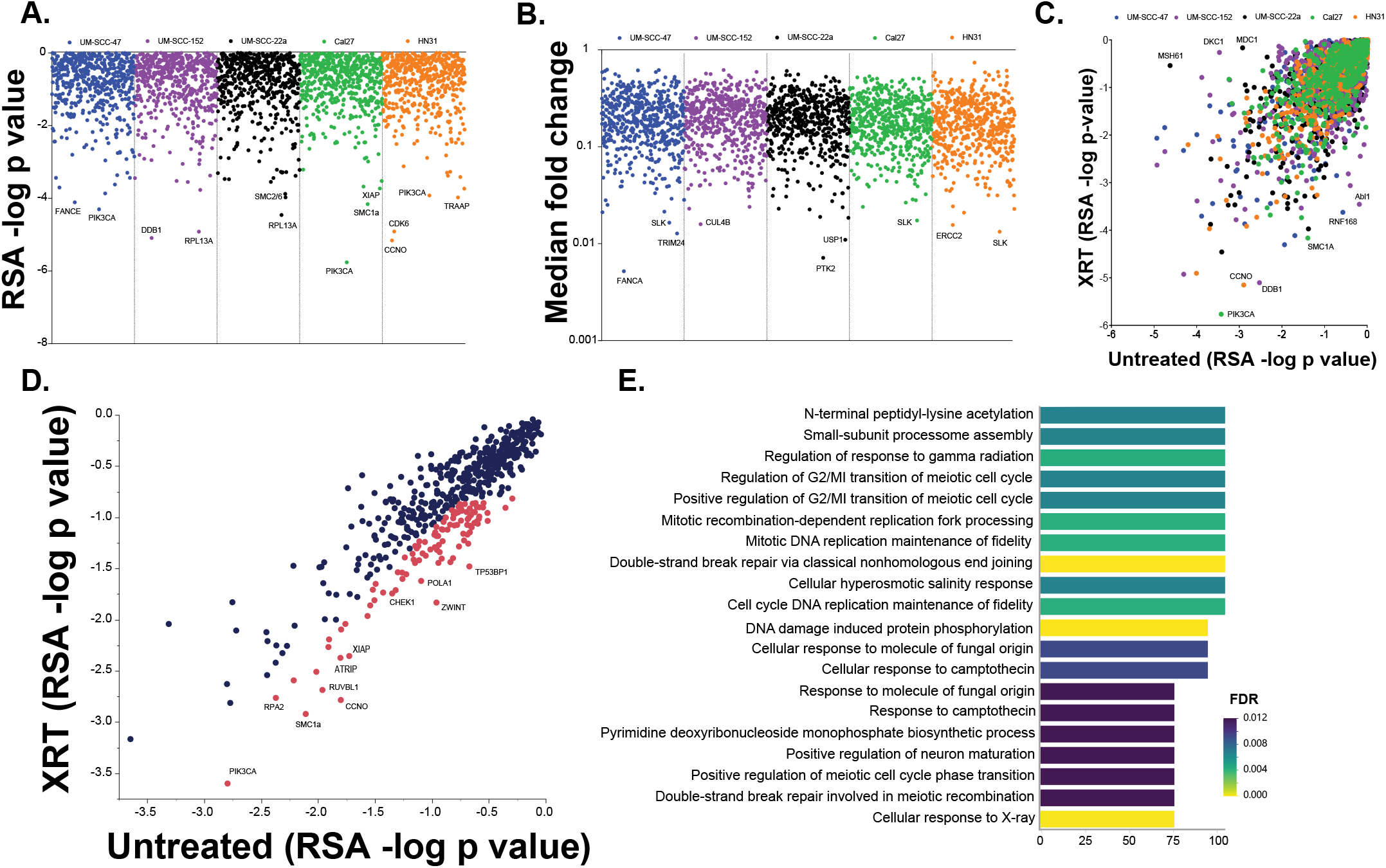
*In vivo* shRNA screening for radiosensitizing targets. A & B) RSA-log p-values (A) and median fold change (B) for each target in our two shRNA screening libraries for each tumor type tested. C & D) RSA-log p-values in irradiated (XRT, y-axis) versus untreated (x-axis) for each tumor type tested (C) and the average for each group (D). Target genes in red were selected for further analysis as potential radiosensitizers and analyzed for Gene Ontology term enrichment (E).

Gene ontology (GO) analysis of targets identified in Fig. 1D was then performed (Fig. E) and, as expected based upon the targets screened, DNA damage repair processes were highly represented in this analysis. However, several additional pathways, particularly protein lysine acetylation, were also identified.

### CREBBP, EP300 or dual specificity protein kinase (TTK) inhibition in combination with radiation in CREBBP/EP300 mutated tumors leads to radiosensitization

In addition to finding general radiosensitization targets, we wished to determine if specific somatic mutations observed in HNSCC were associated with targets, with a goal of identifying genomically-associated radiosensitizers. To accomplish this, we compared the existing RSA log p-values and median fold change data for radiosensitizing targets between tumors that are wild type or mutant for somatic mutations that are represented by the models in the study. This was defined as at least 2 of 5 models harboring a mutation that is observed in >10% of HNSCC. The specific comparisons were: i) CREBBP/EP300 (HN31 & CAL27 vs. UM-SCC-47, UPCI-SCC-152 & UM-SCC-22a), ii) NOTCH (HN31, UM-SCC-47, & UM-SCC-22a vs. CAL27 & UPCI-SCC-152) and iii) CASP8 (UM-SCC-47, UM-SCC-22a & CAL27 vs. HN31 & UPCI-SCC-152) (Fig. 2A-C).

**Fig. 2:**
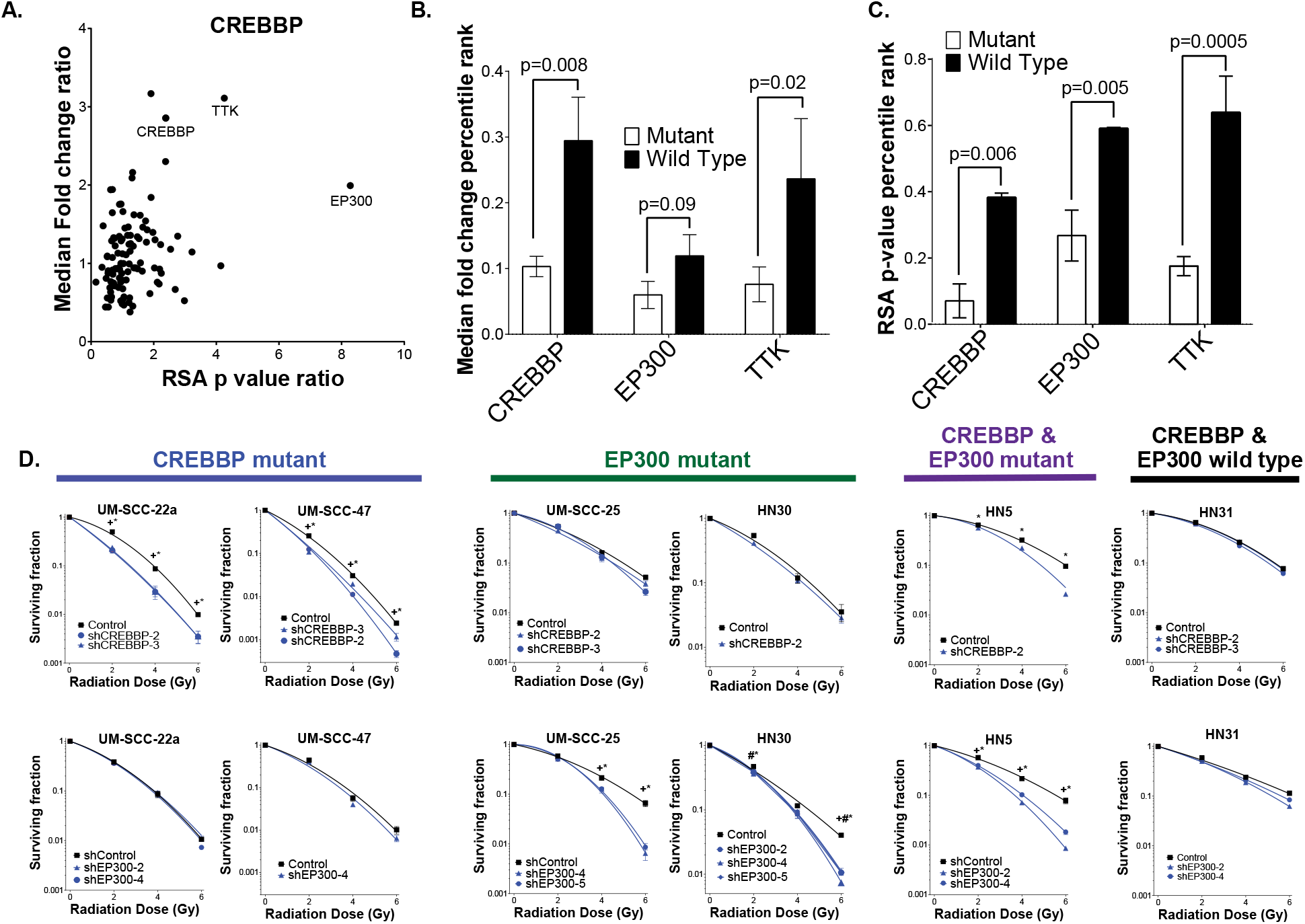
*In vivo* screening identifies genomically-associated radiosensitization in CREBBP/EP300 mutated tumors. A) Ratio of CREBBP mutant vs. wild type for target fold change (y-axis) and RSA log p-value (x-axis) for radiosensitizing targets selected from Fig. 1. B & C) Difference between CREBBP/EP300 mutant and wild type tumors from the *in vivo* shRNA study in as a function of target fold change (B) and RSA log p-value (C). D) Clonogenic assays following irradiation of HNSCC cell lines expressing control and either CREBBP or EP300 shRNA. In B & C, comparisons were evaluated using ANOVA with post-hoc Tukey’s t-test. In D, each point for each group was compared to control using Student’s t-test. For * - shCREBBP-2 and # - shCREBBP-3, p<0.05. All p-values two sided.

While no differences were observed in the comparison CASP8 wild type and mutant tumors in either of the libraries, several targets seemed to preferentially increase sensitivity to radiation in NOTCH and CREBBP mutant tumors (Fig. 2A and Supplementary Fig. 1). Inhibition of Abl1, CCNO and KDM1a appeared to preferentially radiosensitize NOTCH mutant tumors (Supplementary Fig. 1), while inhibition of the CREBBP and EP300 genes, as well as the dual specificity protein kinase (TTK), were associated with increased *in vivo* sensitivity to radiation in the CREBBP mutant tumors in this screen (Fig. 2A).

### Inhibition of CREBBP or EP300 expression leads to in vitro radiosensitization, but only in the presence cognate mutations

Based upon the degree of effect as well as the identification of lysine acetylaton as an enriched pathway in radiosensitizing targets, we chose CREBBP and EP300 for further validation by performing targeted knockdown (KD) across an expanded set of cell lines. We utilized shRNA KD to either CREBBP or EP300 in six HNSCC cell lines (Fig. 2D and Supplementary Fig. 2) of varying CREBBP/EP300 mutation status (see Supplementary Table 1 for details). These cell lines were then treated with radiation, and clonogenic survival was assayed (Fig. 2D). Similar to our *in vivo* screening results, CREBBP or EP300 KD was associated with increased sensitivity to radiation. However, the sensitivity appeared only in the context of a mutation in the cognate gene. For example, CREBBP KD, but not EP300, led to significant radiosensitization in the CREBBP mutant cell line UM-SCC-22a, which is EP300 wild type. Similarly, EP300 KD, but not CREBBP, led to significant radiosensitization in the EP300 mutant cell line UMSCC25 that is wild type for CREBBP. This pattern was consistently observed over all cell lines tested (Fig. 2D).

### CREBBP inhibition leads to increased apoptosis and decreased BRCA1 foci formation following radiation in mutant cells

To further evaluate the observed radiosensitization, we examined apoptosis in multiple cell lines expressing shRNA to CREBBP (Fig. 3A-B). The combination of CREBBP inhibition and radiation led to dramatically increased TUNEL staining in CREBBP mutant (but not wild type) cell lines (Fig. 3A). Similar results were observed on immunoblot examining caspase-3 cleavage (Fig. 3B). We also examined the DNA damage response via immunofluorescence staining of DNA damage foci (Fig. 3C). Importantly, γ-H2AX foci, a marker of DNA damage, were increased following CREBBP KD and radiation in all 3 CREBBP mutant cell lines examined. In contrast, under the same conditions BRCA1 foci induction was significantly reduced in all 3 CREBBP mutant cell lines and HN30 (Fig. 3C). In the HN30 line γ-H2AX induction was actually reduced following radiation and CREBBP KD, likely indicating a different biology occurring in this TP53 and CREBBP WT cell line. Irrespective of mutational status, inhibition of CREBBP generally had little effect on 53BP1 foci formation following radiation (Fig. 3C).

**Fig. 3:**
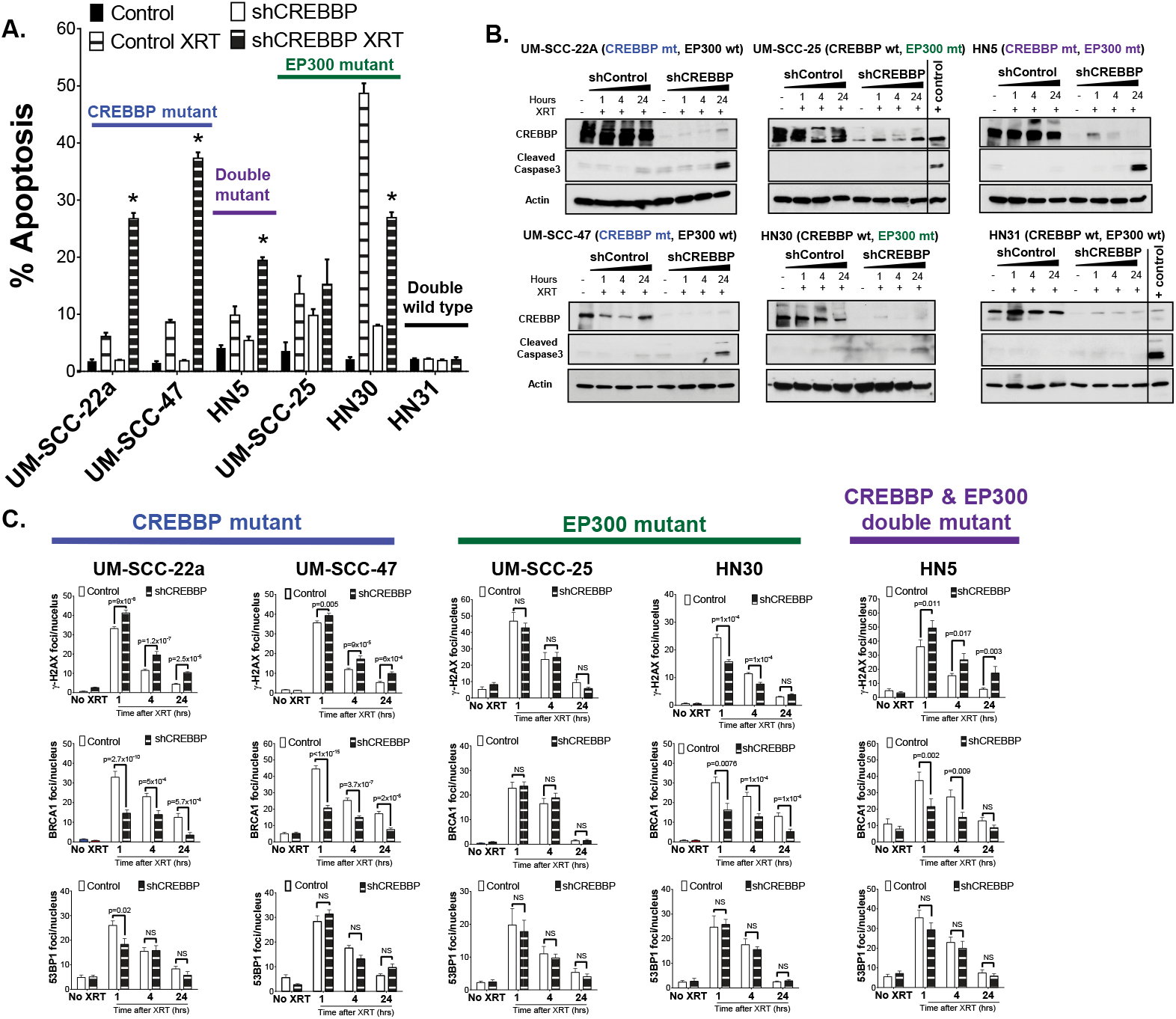
Impaired HR and increased apoptosis is observed following CBP inhibition in CREBBP mutant cells. A & B) Tunel staining (A) and caspase 3 immunoblot (B) following irradiation in HNSCC cells expressing control or CREBBP shRNA. (C) γ-H2AX, 53BP1, and BRCA1 foci following irradiation in shControl or shCREBBP HNSCC cells. Significance tested via students t-test. For A, * indicates a two-sided p<0.05 compared to the Control XRT group. For B & C two sided p-values are indicated.

### Knockdown of CREBBP leads to dramatic in vivo radiosensitization

We further evaluated the therapeutic potential of targeting CREBBP using two separate *in vivo* models of HNSCC. In the first study, we used the CREBBP mutant cell line UM-SCC-47 to generate tumors in the mouse flank. Tumors were treated with 2 Gy x 8 days, in a fractionation scheme designed to recapitulate that used in patients, albeit to a much lower dose (16 Gy total vs. 70 Gy in the clinic). In this experiment, radiation or CREBBP KD (using two distinct shRNA constructs) alone had minimal to modest effect; however, the combination led to a profound tumor growth delay and improved survival (Fig. 4A-B). Indeed, at the conclusion of the tumor growth delay experiment, 5 tumors in the irradiated shCREBBP-2 group (45%) and 7 tumors in the irradiated shCREBBP-3 group (64%) had regressed below the limits of detection. An additional separate experiment was performed to measure apoptosis. TUNEL staining dramatically increased in the combined shCREBBP and radiation groups 8h following the final dose of radiation (2 Gy x 8 d) compared to both the irradiated shControl tumors and unirradiated CREBBP knock down tumors (Fig. 4C). We performed a similar experiment using tumors derived from cells of another CREBBP mutant line, UM-SCC-22a (Fig. 4D). While inhibition of CREBBP alone had a significant effect in this model, we again observed a profound radiosensitization compared to the unirradiated control tumors. In this model, virtually all of the irradiated tumors in both CREBBP knockdown groups regressed below the limits of detection.

**Fig. 4:**
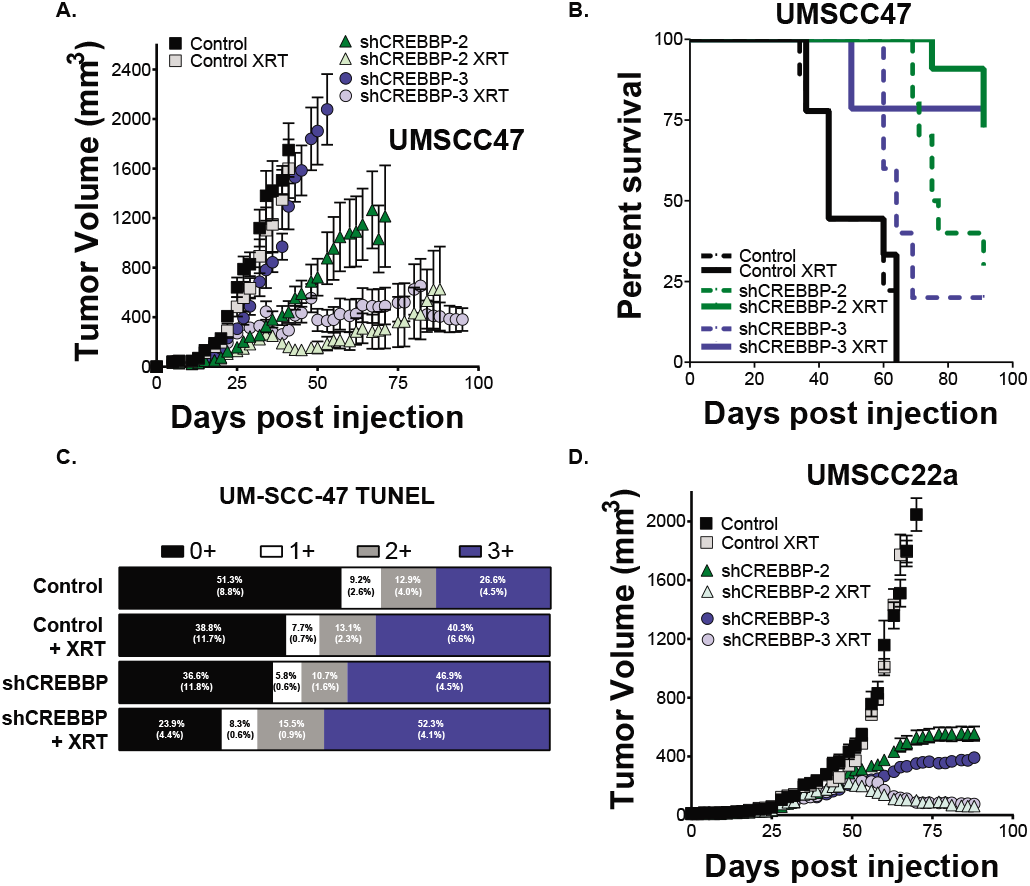
Inhibition of CBP in CREBBP mutant tumors leads to radiosensitization in in two distinct *in vivo* HNSCC models. A & B) Tumor growth delay (A) and survival (B) in a UM-SCC-47 xenograft model expressing either control or two different CREBBP shRNAs following irradiation at 2 Gy/day for 5 days. C) *In vivo* TUNEL staining from UM-SCC-47 xenograft tumors collected 8h following the final dose of radiation in a concurrent experiment with (A). C) A similar experiment performed in the UM-SCC-22a xenograft model. For both models, the slope of the linear regression for both irradiated CREBBP KD groups was significantly different from other groups with at least a twosided p<0.01. In B, log rank statistics comparing shCREBBP-2 XRT or sh-CREBBP-3 XRT groups compared to Control XRT showed a significant increase in survival (twosided p<0.05) in both groups.

### Inhibition of CBP and p300 histone acetyltransferase (HAT), but not bromodomain, function, leads to radiosensitization in CREBBP/EP300 mutants associated with repression of HR

To further examine the therapeutic relevance of the observed radiosensitization in HNSCC, we utilized several chemical inhibitors of CREBBP and/or EP300 function: 1) ICG-001, a CBP specific inhibitor that is thought to inhibit the interaction between CBP and β-catenin, although it is also known to have β-catenin independent effects (note: PRI-724 is an active enantiomer of ICG-001)^19-21^; 2) GNE-272, a bromodomain specific inhibitor for both CREBBP and EP300^22^; 3) A-485, a histone acetyltransferase inhibitor specific for CREBBP and EP300 (note: A-486 is an inactive analog and used as a negative control)^23^. Similar to shRNA-based inhibition of CREBBP, ICG-001 led to significant *in vitro* radiosensitization on clonogenic assay, but only in those cell lines harboring a CREBBP mutation (Fig. 5A-D). This effect was largely due to increased apoptosis following the addition of ICG-001 to radiation (Fig. 5E). Although the analogue of ICG-001, PRI-724, is actively in clinical trial development, neither of these agents target p300, and as predicted, we did not observe sensitization in a wild type or an EP300 mutant cell line (Fig. 5C-D).

**Fig. 5:**
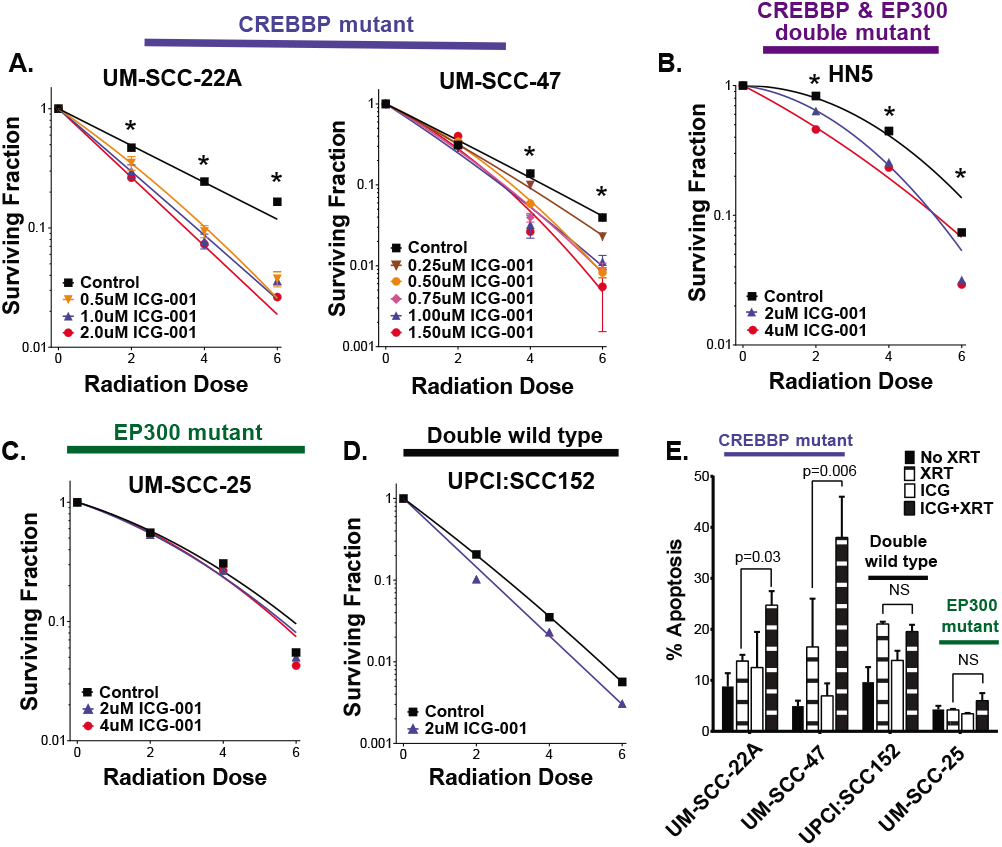
ICG 001 radiosensitizes CREBBP mutant, but not wild type, cell lines. A-D) Clonogenic survival following irradiation and ICG-001 in CREBBP mutant (A & B) and wild type (C & D) cell lines. E) Tunel assay following the same combination (p-values are two-sided and derived from Student’s t-test). Clonogenic survival curves analyzed as in Fig. 2.

Thus, to maximize the clinical impact of the observed radiosensitization, we examined additional inhibitors, currently in clinical development, that inhibit both CBP and p300 function. We initially tested GNE-272, a bromodomain inhibitor; however, this agent had no effects on sensitivity to radiation on clonogenic assay (Supplementary Fig. 3), independent of CREBBP or EP300 status. However, the HAT inhibitor A-485, but not the inactive A-486 analog, led to a profound radiosensitization in cell lines harboring a mutation in either CREBBP or EP300 (Fig. 6A), but not in wild type cell lines (Fig. 6B). The observed radiosensitization was largely due to increased apoptosis following the combination of A-485 and radiation (Fig. 6C-D).

**Fig. 6:**
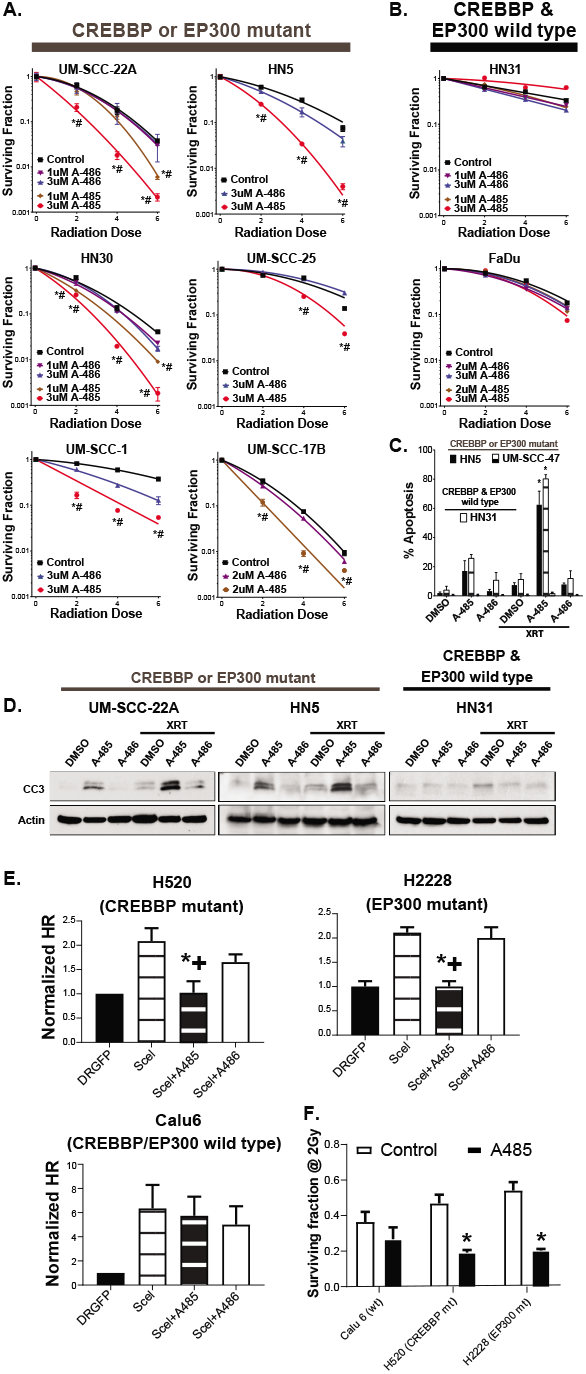
The HAT inhibitor A485 exhibits radiosensitizes cell harboring CREBBP and EP300 mutations. A & B) Clonogenic survival following irradiation and either A485 (active) or A486 (inactive) in CREBBP/EP300 mutant (A) or wild type (B) HNSCC cells. C & D) TUNEL assay (C) and Caspase 3 cleavage (D) in UM-SCC-47 (CREBBP mut), HN5 (CREBBP/EP300 mut) and HN31 (CREBBP/EP300 wt) cells treated with irradiation and either A485 (active) or A486 (inactive). E) iScel assay for HR as described in the methods following treatment with either A485 or A486. F) Survving fraction at 2 Gy from clonogenic assays following exposure to A485. *,+-<0.05 vs. Control (*) and A486 (+). p-values are two-sided and derived from Student’s t-test.

Because of the observed effects of shCREBBP on BRCA1 foci formation following radiation, and the relationship between BRCA1 and homologous recombination (HR), we wished to evaluate the relationship between HAT inhibition and homologous recombination directly via iScel assay^24^. Due to the inability to reliably express the required plasmids in HNSCC cell lines, we utilized lung cancer cell lines which also harbor mutations in either CREBBP or EP300 (Fig. 6E-F). Treatment with A485 led to repression of HR in CREBBP or EP300 mutant cell lines, but not in a wild type line (Fig. 6E). Similar effects were found on clonogenic survival (Fig. 6F), with decreased survival in both mutant lines following exposure to A485 and radiation, but not in the wild type line.

### The observed radiosensitization in CREBBP/EP300 mutants is not CREBBP or EP300 expression level dependent

One potential explanation for the observed radiosensitization is a simple dose-dependency. Namely, if basal levels of CREBBP or EP300 are significantly diminished in mutant cells (and tumors), a more profound inhibition is possible, leading to a more pronounced phenotype. Thus, the observed effect could be due to a more complete inhibition of the protein in the setting of a loss of function mutation. To further explore this hypothesis, we examined basal expression in HNSCC cell lines and tumors in the context of various CREBBP and EP300 mutations. Interestingly, basal CREBBP and EP300 gene expression in the cell lines used in this study were not directly associated with underlying mutation (Supplementary Fig. 4A). Moreover, in a panel of 82 HNSCC cell lines, neither CREBBP nor EP300 mutation was directly associated with mRNA expression^25^ (Supplementary Fig. 4B). Similarly, neither CBP nor p300 protein levels were associated with mutation in the cell lines used in this study (Supplementary Fig. 4C).

In clinical samples from the TCGA HNSCC cohort, no significant difference in CREBBP or EP300 gene expression was observed in mutant tumors (Supplementary Fig. 4D). Moreover, even in the context of nearly complete inhibition of CREBBP protein expression (Fig. 3B & Supplementary Fig. 2) neither HN31 nor HN30 (CREBBP wild type cell lines) were sensitized to radiation (Fig. 2D). Conversely, incomplete inhibition of EP300 (in the case of EP300 mutants HN5 and HN30) led to significant radiosensitization (Fig. 2D & Supplementary Fig. 2).

### Histone acetylation is associated with radiosensitization in CREBBP/EP300 mutants following CBP/p300 targeting in HNSCC

Based on the observed sensitization following HAT, but not bromodomain inhibition, and the lack of evidence for this phenomenon being solely gene expression dependent, we further investigated histone acetylation status in HNSCC cell lines following combination treatment. As expected, treatment with A-485 inhibited histone acetylation at H3K18 and H3K27 in HN5 and UM-SCC-22a cells (Fig. 7A). Both cell lines harbor mutations in either CREBBP and/or EP300 and exhibit sensitivity when HAT inhibition is combined with radiation. Interestingly, FaDu, a CREBBP/EP300 wild type cell line with no observed radiosensitization, showed no effect on histone acetylation following treatment with HAT inhibitor at baseline or following radiation treatment. Similar effects on histone acetylation were observed following the expression of shRNA specific to CREBBP (Fig. 7B) and treatment with ICG-001 (Fig. 7C). *Selected mutations in CREBBP exhibit increased acetylation activity consistent with gain of function and potentially mediating response to radiation*.

**Fig. 7:**
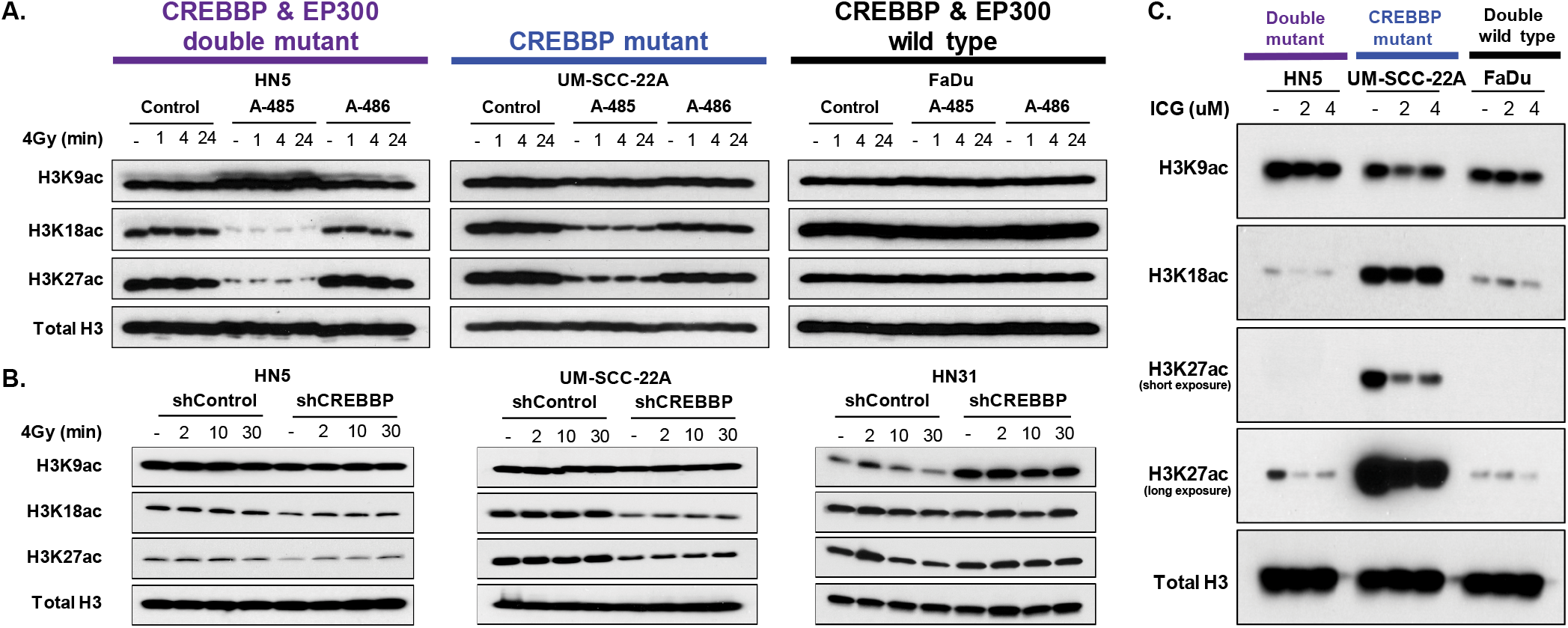
Cells with mutations in CREBBP exhibit inhibition of histone acetylation following inhibition of CBP function. A-C) Histone acid extraction shows that A-485 (A), shCREBBP (B) and ICG-001 (C) leads to inhibition of H3K18ac and H3K27ac in cell lines that harbor a mutation in CREBBP or EP300, but not those that are wild type for both.

Previously, mutations within the inhibitory TAZ domain of CBP have been linked to increased histone acetylation^26^. Because of this observation, we evaluated two of our cell lines (UM-SCC-22a and UM-SCC-17b) with similar mutations (Fig. 8A-B). Compared to wild type cells, these cell lines exhibited profoundly higher levels of acetylation of several histone marks as well as increased global protein acetylation (Fig. 8A). Additionally, CBP protein itself was acetylated at higher levels compared to wild type cells. As expected, the HAT inhibitor A485 inhibited the observed acetylation in both mutant cell lines (Fig. 8B).

**Fig. 8:**
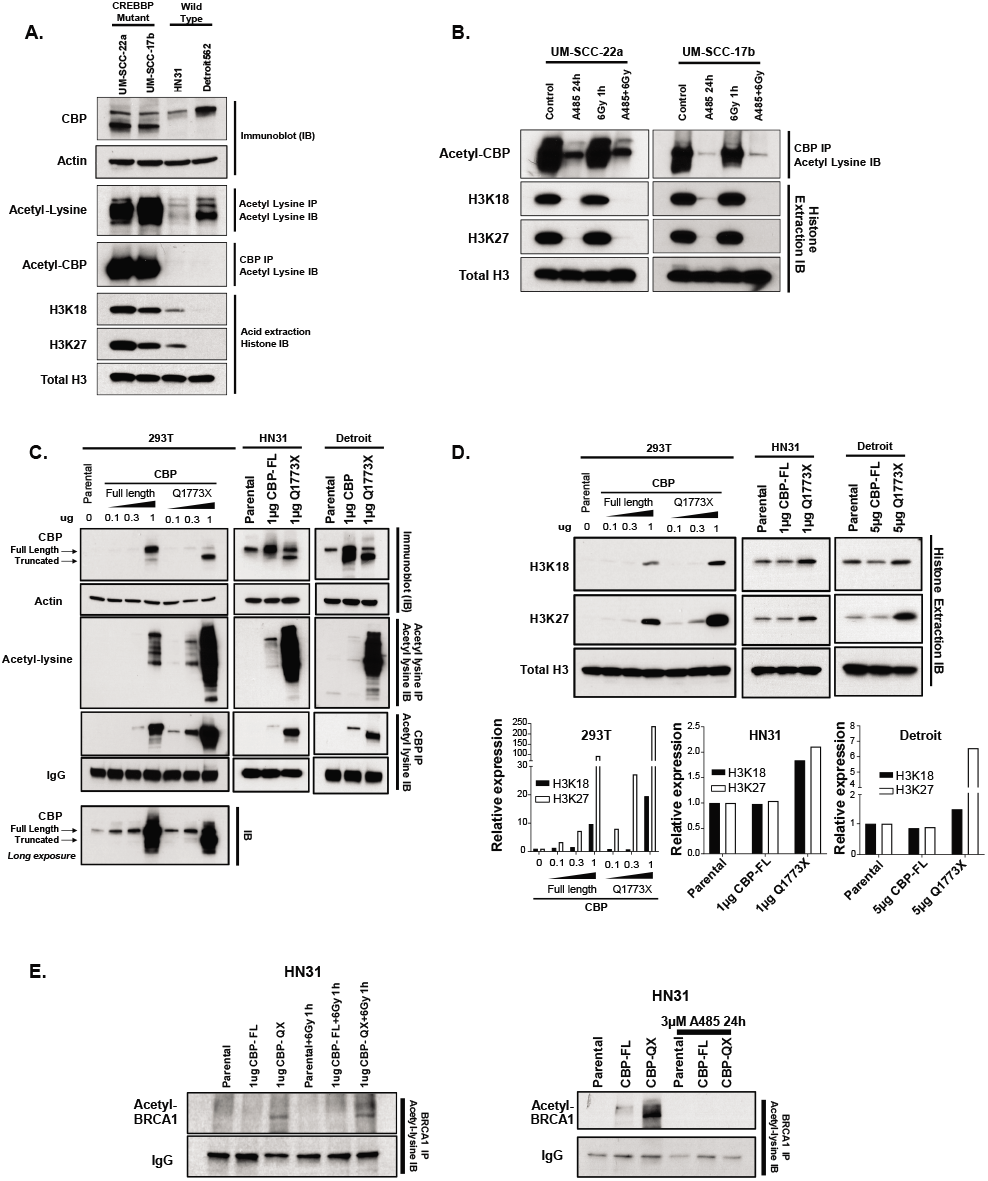
Gain of function mutation in CREBBP mutant cells. A-E) Immunoblot (IB), histone acid extraction or immunoprecipitation (IP) for either CBP, acetyl-lysine or BRCA1 was performed as indicated in the individual panels. A) Cell lines with a CREBBP mutation in the TAZ2 domain have increased global protein acetylation (IP acetyl lysine & IB for acetyl lysine), CBP-auto acetylation (IP CBP & IB for acetyl lysine) and histone acetylation (acid extraction and IB) compared to wild type HNSCC cell lines. B) CBP-auto and histone acetylation are inhibited by A485 in CREBBP mutant lines. C & D) Forced expression of TAZ mutant CBP (Q1773X) in 293T, HN31 and Detroit cell lines leads to increased global protein and CBP-auto acetylation (C) as well as histone (D) acetylation (densitometry below blot), compared to forced expression of wild type. E) Forced expression of CBP mutant leads to potential increase in BRCA1 acetylation, which is slightly increased by radiation and inhibited by A485.

We then forced CREBBP wild type (293T and HN31) cells to express full length or a representative truncated inhibitory region mutant (Q1773X). In both lines, expression of the mutant led to a profound increase in CBP auto-acetylation and global protein lysine-acetylation that was not observed following forced expression of full length CBP (Fig. 8C). Similarly, forced expression of mutant CBP led to increased acetylation at H3K18 and H3K27 histone marks (Fig. 8D). Additionally, we specifically examined the effects of mutant CBP on BRCA1 acetylation, which was similarly increased following forced expression of mutant CBP compared to wild type (Fig. 8E). *Mutations in CREBBP/EP300 are associated with outcome following radiation in SCC*.

To examine the clinical association between CREBBP/EP300 mutation – and other mutations in HNSCC – with radioresponse, we identified a cohort of patients within the Head and Neck Cancer Genome Atlas (TCGA) that were denoted as having radiation as a component of their therapy (Supplementary Table 4). Genes mutated at ≥10% frequency (high enough for potential clinical utility) were then queried to determine their relationship to overall survival (Fig. 9A). Only 3 genes (TP53 (p=0.015), CASP8 (p=0.055) and CREBBP/EP300 (p=0.046) were associated with OS, as was the presence of HPV (p=0.002) (47 patients, 17.4%) (Fig. 9A). Tumor stage (p=0.69), nodal stage (p=0.61), and tumor site (p=0.79) were not significantly associated with survival in this population; this is likely due to the similar clinical characteristics of the selected cohort, all of whom had advanced stage disease treated with combined modality therapy. While TP53 was associated with OS in patients who did not receive radiation (HR 1.52, p=0.053), CASP8 (p=0.41) and CREBBP/EP300 (p=0.89) were not, indicating that in the latter genes this phenomenon may be dependent upon radiation response.

**Fig. 9:**
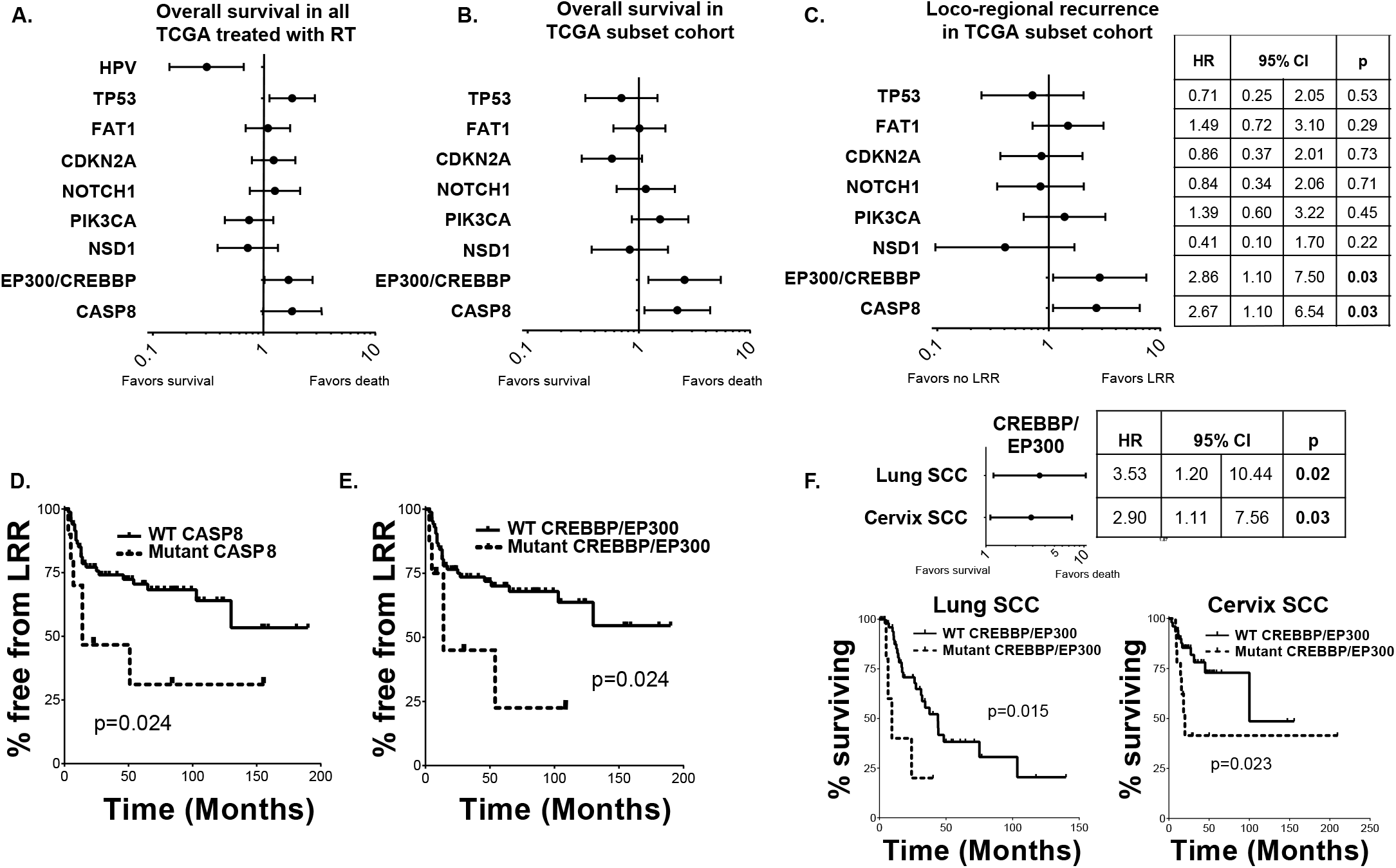
Whole exome sequencing identifies CREBBP/EP300 mutation significantly associated with outcome and treatment failure in several SCC cohorts. A) Forrest plot of hazard ratios (HR) for Overall survival (OS) for TCGA patients known to have received radiation therapy. B & C) Forrest plot of HR for OS (B) & loco-regional recurrence (LRR) (C) in a subset of TCGA patients treated uniformly with surgery and radiation and known patterns of failure. B & C) Kaplan-Meier curves for LRR in patients with either CASP8 (D) or CREBBP/EP300 (E) mutations. F) Forrest plot and KM curves for OS by CREBBP/EP300 mutation in Lung and Cervix SCC cohorts.

Because the TCGA includes a cohort of patients with heterogeneous treatments and without details of patterns of failure (particularly the difference between LRR and DM), we examined patient outcomes in a more uniformly treated cohort of patients within this group, in which treatment and patterns of failure details were enumerated (Fig. 9 B-C). This subset included 94 patients with HPV-negative HNSCC treated uniformly with surgery and post-operative radiation (Clinical characteristics in Supplementary Table 5). In this analysis, TP53 (as a binary variable), was not associated with OS or LRR (Fig. 9B-C). Mutations in both CASP8 and CREBBP/EP300 were associated with significantly reduced overall survival and higher rates of LRR in this patient population (Fig. 9B-E).

Since CREBBP/EP300 are commonly mutated in squamous tumors^27^, we expanded our analysis to other SCCs to determine whether their mutation might also be associated with poor outcome in other SCC tumor types treated with radiation. Among the TCGA squamous tumors, both lung and cervix had sufficient numbers and treatment annotations for analysis (Clinical characteristics in Supplementary Table 6). We found that CREBBP/EP300 mutations were associated with poorer overall survival in radiation treated patients with lung and cervix squamous tumors (Fig. 9F).

## Discussion

There are no biologically driven precision medicine approaches to radiation therapy, with treatment largely guided by clinical stage and intensified via the addition of cytotoxic chemotherapy. This leads to high degrees of toxicity as well as both over- and undertreatment depending upon the patient and tumor. HNSCC is no exception to this phenomenon, with a highly toxic standard treatment of concurrent chemoradiation that has largely remained unchanged for decades. To improve this paradigm, we performed *in vivo* screening of HNSCC models and identified both general radiosensitizing targets as well as a novel genomically-associated sensitization. This latter effect is potentially related to a gain of function in mutant CBP and p300 leading to increased basal acetylation and BRCA1 function, rendering these tumors highly sensitive to the combination of HAT inhibition and radiation (potential mechanism in Fig. 10).

**Fig. 10:**
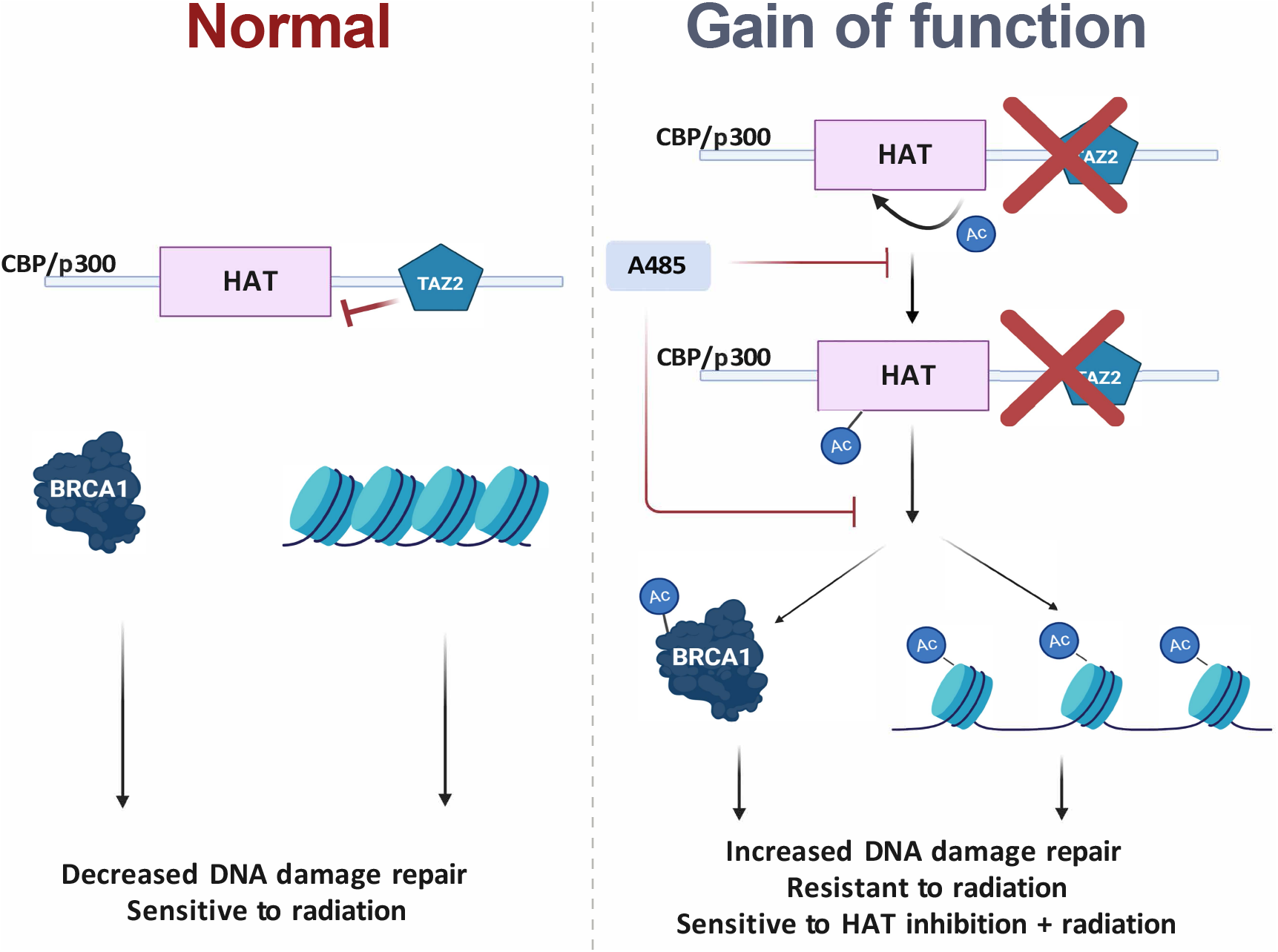
Proposed mechanism CREBBP/EP300 mutant specific radiosensitization.

This work is the first, to our knowledge, to perform an *in vivo* shRNA screen for targets associated with radiosensitization. We choose HNSCC as our initial model, both due to the primacy of radiation in curative therapy as well as the relative dearth of targetable genetic alterations in this malignancy. Importantly, the use of *in vivo* screening takes into account both tumor bulk, metabolism, angiogenesis and stromal interactions which are not identified in *in vitro* screens. The identification of several genes identified as targets for clinical radiosensitization in HNSCC by our own group and others – notably CHEK1^28,29^, PIK3CA^15,30^, PTK2^16^ and XIAP^14^ – argues for the utility of this technique in identifying more relevant targets for clinical trial development. Validation of additional general targets for radiosensitization identified in a similar manner is ongoing.

However, despite the potential for these targets to improve response, toxicity remains a concern for agents, such as PI3K^31,32^ or CHK1/2 inhibitors^33^, which exert broad anti-tumor effects when they are combined DNA damage-based therapies in the clinic. One means of partially mitigating the overlapping toxicities of radiation and targeted therapies is tailoring of specific agents to genomic events that drive radioresistance. Evaluating our screening data in this manner we found that inhibiting the protein acetylation function of CBP and p300 led to a dramatic sensitization to radiation in CREBBP/EP300 mutant tumors and cell lines.

The concept of genomically-driven radiation targeting is largely in its infancy.

Despite ample evidence of genomic dependencies for targeted therapies – the classical example being PARP inhibition in BRCA-altered tumors^12^ – direct links between a particular somatic mutation or genetic alteration and sensitivity to the combination of radiation and a particular agent are limited. Indeed, most studies of radiosensitizers have focused broadly on agents that either effect DDR or inhibit kinases known to be important in tumorigenesis. This approach has the advantage of a potentially broad applicability but risks masking effects in specific groups of patients ultimately leading to underperforming clinical trials and/or unacceptable toxicities. Conversely, identifying a specific effect in CREBBP/EP300 mutants, using a HAT inhibitor being developed for clinical use and radiation, can maximize response and tailor therapy to achieve a minimum of toxicity.

The identification of genomically-associated radiosensitization in CREBBP/EP300 mutants is particularly of interest in HNSCC, and indeed in SCCs in general, as we have shown in the current study that mutations in these genes are associated with clinical radioresistance. This is the first large scale examination of somatic mutations in this context and serves to link particularly treatment resistant tumors with a genomically-tailored therapy.

CREBBP and EP300 are homologous multifunctional bromodomain-containing acetyltransferases. Although these genes are mutated in HNSCC, they have not been extensively studied in this tumor type. Specifically, CREBBP and EP300 are collectively mutated in 13% of HNSCC ^34^, with similar frequencies in both HPV-positive and HPV-negative disease. Missense mutations are clustered in the acetyltransferase domain, and there is a reasonable frequency of truncating mutations (~20%). Additionally, many of these mutations are heterozygous, indicating possible haploinsufficiency, as has been seen for the chromatin modifying genes in the BAF complex ^35^. CREBBP and EP300 are also mutated in 14% of all squamous cancers^27^, and we found them associated with poor survival in lung and cervical tumors (Fig. 9F). Although much of our work has been done in HNSCC, it should be applicable to other tumor types, particularly squamous cell carcinoma. Furthermore, it is possible that the mechanism can be expanded to include other acetylation or chromatin modifying genes and shed new light on the role of acetylation in the DNA damage response and repair.

Previously one study has identified an “addiction” to p300 in the context of CBP deletion, which was felt to replicate naturally occurring mutations in CREBBP ^36^. However, in our study we did not observe an effect with p300 inhibition in CREBBP mutant cell lines in our model, either at baseline or in combination with radiation. We also did not observe the converse in EP300 cell lines when CBP was inhibited. Thus, deletion of either protein may not recapitulate naturally occurring mutation.

Indeed, the data support a potentially more complex hypothesis of a gain of function for at least some mutations in CREBBP/EP300. We observed high basal levels of both protein, including BRCA1, and histone acetylation in cell lines harboring mutations which truncate the protein downstream the HAT domain, with forced expression of similar mutations in wild type cell lines recapitulating this effect. This acetylation is reversed following inhibition of CBP and p300 HAT activity.

A gain of function for CBP and p300 is not wholly unprecedented, as previous data from the Cancer Cell Line Encyclopedia suggested a gain of function for certain CREBBP mutants, specifically truncation mutations located in the TAZ domain, for acetylation at certain histone marks, although this was only identified in a few cell lines^26^. However, based on our data, this gain of function – albeit with varying degrees of basal activity – may be more prevalent than previously appreciated. Moreover, this gain of function appears to extend to both increased auto-acetylation as well as acetylation of additional proteins, notably BRCA1. This would be in addition to a basal increase in acetylation of histone tails (H3K18, H3K27, H4K5/8/12/16), which serves to allow DDR proteins to access damaged DNA and facilitate its repair more easily^37,38^. We believe that this state functions to generally promote a state primed to repair DNA via increased BRCA1 activity and HR, compared to cell lines lacking these mutations. This state, in turn, renders the combination of HAT inhibition and radiation highly lethal, via an inhibition of HR (see Fig. 10).

This study is limited in that the full spectrum of mutations in CREBBP and EP300 have not been studied. It is possible that the observed sensitization could be due to a combination of factors related to both a gain and loss of function for these proteins, particularly as the effect of missense mutations in the HAT domain, which are a significant proportion of all CREBBP/EP300 mutations, is unclear. Similarly, we are not certain about the most appropriate terminology to use for this phenomenon. Our screen was analyzed to identify genomically-associated sensitization, with several targets identified. However, the relationship between CREBBP/EP300 mutations and genomically-dependent targeting data is something akin to context-dependent oncogene addiction. However, many CREBBP/EP300 mutations have patterns consistent with loss of function, so more mechanistic studies are necessary to clarify the phenotype. Additionally, although we have identified BRCA1 and HR as likely mediators of this phenomenon, because of the relatively broad effects of the examined mutations on protein acetylation, additional studies are needed to determine if additional signaling pathways modulate this effect.

In conclusion, we have both identified prognostic markers of outcome following radiation in SCC as well as explored a novel radiosensitization involving one of these biomarkers, mutated CREBBP/EP300. This genomically-associated radiosensitization appears to specifically involve effects on DNA damage repair, leading to a HR deficiency following DNA damage and leading to increased apoptosis. A gain of function effect in mutated cell lines may be driving this phenomenon leading to a basal hyperacetylated state affecting BRCA1 function, which is inhibited using a HAT inhibitor. This agent is currently being explored for clinical trial use, and thus could be translated clinically to improve outcomes in SCC.

## Materials and Methods

### Cell Lines

HNSCC cell lines (UM-SCC-47, UM-SCC-22a, UM-SCC-25, UM-SCC-1, HN31, HN30, UM-SCC-17b, UPCI:SCC152, Cal-27, UD-SCC-2 and HN5) used in this study were generously supplied by Dr. Jeffrey Myers via The University of Texas MD

Anderson Cancer Center Head and Neck cell line repository. HEK-293T, NCI-H520, NCI-H2228, Calu-6, FaDu and Detroit562 were purchased form American Type Culture Collection (Manassas, VA). Cell lines were tested for mycoplasma and genotyped before experiments. See the Supplementary Methods for more details.

### In vivo shRNA screen

Each library was cloned into the pRSI17 vector (Cellecta) and packed into lentivirus particles. HNSCC cell lines were infected *in vitro* through spinfection with virus containing the library at a low MOI (~20% infected cells as measured by flow cytometry) in order to minimize superinfection of cells. Cells were selected with puromycin for at least 2 days and grown *in vitro* for <3 population doublings prior to injection of 4 million cells subcutaneously in nude mouse flank. An additional 2 million cells from the day of injection were collected as a frozen reference cell pellet. Pilot studies were performed to: i) examine the frequency of tumor initiating cells (TIC) and determine whether the cell line could maintain shRNA library complexity *in vivo* and ii) identify the dose of radiation needed to achieve ~20% tumor reduction for each model by the conclusion of the experiment.

For the screening experiment itself, xenografts were treated with 2 Gy/day of radiation once the tumor had reached approximately 100 mm^3^ to a total dose of 6-10 Gy depending upon the model. Following treatment, the tumors were allowed to grow for approximately 2 weeks (volume ~ 500mm3). DNA was isolated from tumor and reference cells, amplified, and sequenced on Illumina sequencers as previously described^39^.

Hairpin counts were normalized to counts per million (CPM) per sample to enable comparison across samples. For each sample, (log2) fold-change of each hairpin in the tumor was calculated compared to the level in the reference pellet. A hairpin summary measure per cell line was derived from the median of quantile transformed log2 FC across replicates. Next, a modified version of the redundant siRNA activity (RSA) algorithm^40^ was used to derive a gene level summary measure per cell line. RSA attempts to provide a gene level summary estimate of the impact of knock-out of the gene by calculating a stepwise hypergeometric test for each hairpin in a gene. Similar to GSEA, it is based on evidence from multiple hairpins of a gene showing an impact of cellular fitness. Our modifications were to ensure both that at least 2 hairpins were used when calculating the minimum p-value (in RSA) and that hairpins ranking above luciferase controls were not used when determining the minimum p-value. Quantile rank of luciferase controls barcodes was determined through evaluation across all experiments; on an average luciferase barcodes ranked >0.6 on the quantile transformed log2fc scale, so hairpins with quantile transformed log2fc > 0.6 were not used for the gene-level RSA score.

### TUNEL Assay

Following experimental treatments, all cells were collected including floating cells and TUNEL staining was performed using the APO-DIRECT Kit (BD Pharmingen) according to the manufacturer’s protocol. Briefly, 1 million cells were fixed in 1% paraformaldehyde on ice for 30min. Cells were then washed in PBS and fixed in 70% ethanol overnight at −20C. Cells were washed twice with provided buffer then stained with 50ul of DNA labeling solution at 37C for 45min. Cells were then rinsed twice with provided buffer and resuspended in 300ul of rinse buffer. Cells were then analyzed by flow cytometry using the BD Accuri C6 flow cytometer (BD Biosciences) with 488nm laser, 533/30 filter and FL1 detector. 10,000 events were measured per sample.

### Histone Extraction

Histone proteins were extracted from treated or untreated cells using a histone extraction kit (Abcam) according to the manufacturer’s protocol. Briefly, cells were harvested, and pellet was obtained by centrifugation at 10,000 rpm for 5 minutes at 4°C. The cells were re-suspended in 1X pre-lysis buffer and incubated at 4°C for 10 minutes on a rotator and then centrifuged for one minute at 10,000 rpm at 4°C. The cell pellet was re-suspended in lysis buffer at a concentration of 200 uL/10^7^ cells and incubated on ice for 30 minutes, then centrifuged at 12,000 rpm for 5 minutes at 4°C. The supernatant was collected and 300uL balance buffer-DTT was added per 1mL supernatant. The quantity of protein extracted was measured with a DC protein assay kit (Bio-Rad, Hercules, CA, USA). 2-4 μg of protein were separated by western blot analysis as described previously.

### Immunofluorescence Staining

Immunofluorescence was performed to measure quantitative differences in DNA damage repair and response. Cells were cultivated on cover slips placed in 35-mm cell culture dishes. At specified time points after exposure to radiation (2Gy), cells were fixed in 4% paraformaldehyde for 10 min at room temperature on shaker, briefly washed in phosphate-buffered saline or PBS (Biorad), and placed in 70% ethanol at 4°C overnight. These fixed cells were washed with PBS twice to remove ethanol and permeabilized with 0.1% IGEPAL (octylphenoxypolyethoxyethanol) for 20 min at room temperature on shaker, followed by blocking in 2% bovine serum albumin (Sigma) for 60 min, and then incubated with anti-γH2AX (1:400, Trevigen), 53BP1 (1:200, CST) or anti-BRCA1 primary antibody (1:500, Santa Cruz) overnight at 4°C. Next day fixed cells were washed three times with PBS and incubated for 45 minutes in the dark in secondary anti-mouse antibody conjugated to FITC to visualize γH2AX or BRCA1. Secondary anti-rabbit antibody conjugated to Cy3 was used to visualize 53BP1. DNA was stained with 4’, 6-diamidino-2-phenylindole (Sigma) at 1:1000 (1 microgram/ml). Immunoreactions were visualized with an Olympus or Leica Microsystems microscope (Wetzlar, Germany), and foci were counted with Image J software (https://imagej.nih.gov/ij/).

### Plasmids and shRNA Transfection

Packaging cell line HEK-293T was co-transfected with 3 microgram GIPZ lentiviral shRNAs specific for the CREBBP, EP300 gene or control(GE Dharmacon) and lentiviral vectors DR8.2 and VSVG (Addgene). Two and three days after transfection, virus containing media was filtered through a 0.45 micron PVDF syringe filter and polybrene was added (1:2000). Target cells were transduced with virus for 4-6h and were subjected to puromycin antibiotic selection. Pooled knockdown cells and counterparts shControl cells were assessed for CREBBP protein expression by immunoblotting. shRNA sequences are given as follow:

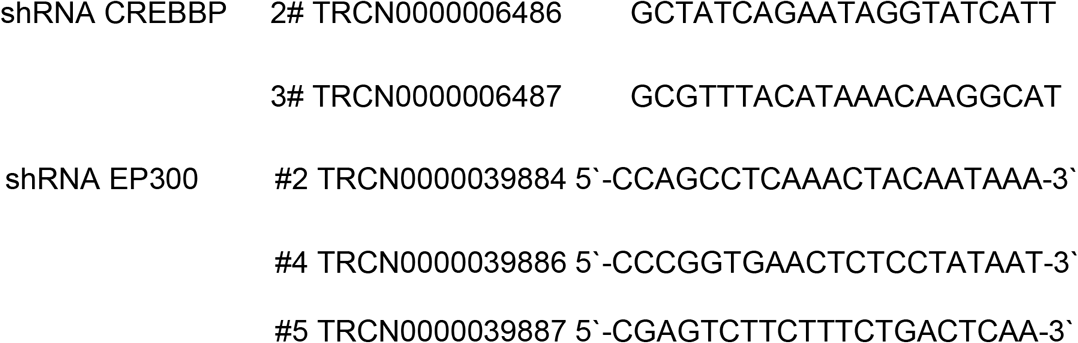

### Site-directed mutagenesis

Site-directed mutagenesis was performed using 50ng of KAT3A / CBP (CREBBP) (NM_004380) Human Tagged ORF Clone (OriGene) as the dsDNA template. This was carried out using the QuickChange II XL Site-Directed Mutagenesis Kit (Agilent Technologies) according to the manufacturer’s protocol with the following exception: One Shot™ Stbl3™ Chemically Competent E. coli (Invitrogen) was used for transformation rather than the XL10-Gold Ultracompetent Cells supplied with the kit because of the dependency on chloramphenicol selection already found in the full length CREBBP vector. Mutagenic oligonucleotide primers were designed using Agilent QuickChange Primer Design program and purchased from Sigma-Aldrich with PAGE purification with the following sequences:

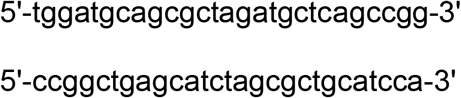

Following transformation, single colonies were selected on LB plates containing 34ug/mL Chloramphenicol, expanded in LB broth containing 34ug/mL Chloramphenicol overnight, and extracted with a QIAfilter midi kit (Qiagen).Sanger sequencing was utilized to confirm the presence of the desired mutation. Mutant plasmids were directly transfected into cell lines using GeneJet transfection reagent (SignaGen Labs) or packaged in 293T for lentiviral infection.

### HR specific DRGFP/DNA repair assay

Calu6 and H520 cells were transfected with 3ug DRGFP substrate using PEI. H2228 cells were electroporated with 3ug DRGFP substrate using reagent T (Amaxa Cell Line Nucleofector Kit T, Lonza), program X-001 on the Nucleofector 2b (Lonza). Puromycin selection was performed with 2ug/ml for Calu6 and H2228 or 4ug/ml for H520. DRGFP expressing cells were electroporated in reagent T for H520 and H2228 or reagent L (Amaxa Cell Line Nucleofector Kit L, Lonza) for Calu6 with 10ug I-SceI using program X-005 for Calu6, T-020 for H520 or X-001 for H2228. Following electroporation, cells were washed with media and transferred into 6 well dishes with 2ml media/well containing either DMSO, 3uM A485 or 3uM A486 and incubated for 48h. Cells were then trypsinized, washed with PBS and resuspended in 300ul of PBS containing 1% FBS, and flow cytometry was run to detect GFP using the BD Accuri C6 flow cytometer (BD Biosciences).

### Mouse xenograft model

In vivo studies were performed following Institutional Care and Use Committee (IACUC) approval. Male athymic nude mice (6-8 weeks old, ENVIGO/HARLAN, USA) were randomly assigned to one of 6 treatment groups of 15 mice each for each cell line tested. Tumor cells (2 × 10^6^ in 0.1 mL of serum-free medium) were injected subcutaneously in the right dorsal flank of each mouse. After palpable tumors had developed, tumor diameters were measured with digital calipers, and tumor volume was calculated as A × B2 × 0.5, where A represents the largest diameter and B the smallest diameter. When the tumor volumes reached approximately ~150 mm^3^, tumors were irradiated with 16 Gy (2 Gy/day x 8 days) and tracked for approximately 4 weeks for tumor growth delay experiments (n=~10/group). At that time, the experiment was completed and tumors harvested. For in vivo TUNEL assay, tumors were collected 8 h after the last radiation treatment (n=5/group). A tumor growth delay experiment was performed in a similar fashion using UM-SCC-22A shControl, UM-SCC-22A shCREBBP#2, and UM-SCC-22A shCREBBP#3. To compare tumor growth delay between groups, linear regression of the growth curve was calculated for each group and the slope of each curve was compared between groups using Graph Pad Prism (v8.0).

### *In vivo* TUNEL assay

Paraffin embedded sections (4μm) of UMSCC47 tumor xenografts were mounted on coated slides and sent to HistoWiz Inc. (histowiz.com) for TUNEL staining and quantification. TUNEL staining was performed using a Standard Operating Procedure and fully automated workflow with Deadend colorimetric TUNEL system from Promega. After staining, sections were dehydrated and film coverslipped using a TissueTek-Prisma and Coverslipper (Sakura). Whole slide scanning (40x) was performed on an Aperio AT2 (Leica Biosystems). Images were analyzed using Halo (version 2.3.2089.34) image analysis software from Indica Labs (Albuquerque, NM). Regions of interest were selected. TUNEL staining was segmented using the CytoNuclear algorithm. Total cell counts were thresholded into low, medium, and high intensity staining bins.

### Clinical data

This study was approved via appropriate Institutional Review Boards. The initial patient cohort consisted of the Head and Neck TCGA group that satisfied the following criteria: i) whole exome sequencing data is available and ii) were denoted in the TCGA records as having received radiation as part of their initial therapy (Clinical characteristics in Supplementary Table 4). Of the 523 patients in the TCGA cohort, a total of 276 patients met these criteria. Whole exome sequencing from these tumors were examined for genes with mutations in ≥10% of tumors and significance on MutSig with the following genes meeting these criteria: TP53, FAT1, CDKN2A, NOTCH1, NSD1, CREBBP and EP300 (combined due to significant homology) and CASP8 ^34,41^. A subset (n=94) of patients from this cohort have known specific treatment characteristics and patterns of failure analysis (Clinical characteristics in Supplementary Table 5). These patients were all treated with surgery and post-operative radiation with well-annotated outcomes, including loco-regional recurrence and distant metastasis. In addition to the above, all tumors in this subset cohort were HPV/p16-negative as demonstrated by immunohistochemistry (IHC) or in-situ hybridization (ISH).

Two additional cohorts from TCGA, the lung SCC and cervical SCC cohorts, were also examined, with a total of 61 and 66 patients respectively annotated as having received external beam radiation and examined for outcomes (Clinical characteristics in Supplementary Table 6).

For all clinical data overall survival was defined from time of diagnosis until death or last follow up. Time to loco-regional recurrence (LRR) or distant metastasis (DM) was defined as time from diagnosis until either an event or last follow up. Univariate analysis was performed using Cox-regression (SPSS v25). Kaplan Meier curves were generated, and group comparisons were performed using log-rank statistics.

## Contributions

HS and CP directed the experiments and analyzed the clinical data. MF assisted in the planning, direction and analysis of the in vivo screening experiments. MK, TX, LY, TH and SS directly performed the *in vivo* screening and/or processed and analyzed the resultant samples. MK, KB, JM, AH and DM generated the stable KD cell lines. DM, JM, KB, MG, AS, AH and MK performed the in vitro experiments. FJ, JW and LS generated and/or analyzed the HNSCC cell line RNA Seq and mutation data. BB, JM and RF reviewed and evaluated the clinical data analysis. MK initiated this study, which was then continued by DM. This determined co-first author order.

