## Supplementary Methods for "Inhibition of histone acetyltranserase function radiosensitizes CREBBP/EP300 mutants via repression of homologous recombination, potentially targeting a novel gain of function"

**Cell lines**

UM-SCC-47, Cal-27 and UM-SCC-25 were maintained in Dulbecco modified Eagle medium (Gibco, USA), supplemented with 10% fetal bovine serum, 1% penicillin/streptomycin, 1% sodium pyruvate, 1% nonessential amino acids, and 2% vitamins. HEK-293T, HN5, HN30, HN31, UM-SCC-1, UM-SCC-17b and UM-SCC-22a were maintained in DMEM/F-12 50/50 medium supplemented with 10% fetal bovine serum and 1% penicillin/streptomycin. UPCI:SCC152, Detroit562, Calu-6 and FaDu were maintained in MEM medium supplemented with 10% fetal bovine serum, 1% nonessential amino acids and 1% penicillin/streptomycin. UD-SCC-2, NCI-H520 and NCI-H2228 were maintained in RPMI 1640 medium with 10% fetal bovine serum and 1% penicillin/streptomycin. All cell lines were incubated at 37^o^C and 5% CO2 atmosphere.

**Clonogenic Survival Assay**

Single cells were plated in 6-well plates overnight. The next day cells were incubated with specified drugs before irradiating at the indicated doses. The cells formed colonies over a 10- to 14-day incubation period andcolonies were fixed in a 0.25% crystal violet/methanol solution. Colonies containing more than 50 cells each were counted. Survival curves were generated using GraphPad Prism.

**Western Blot Analysis**

Cells were washed in PBS and scraped and collected in sufficient amount of whole cell lysis buffer (20mM HEPES pH 7.9, 0.4M NaCl, 0.1mM EDTA pH 8, 0.1mM EGTA, pH 7, 1% Igepal, 1X Halt protease inhibitor cocktail and 1X Halt phosphatase inhibitor cocktail (Thermo Sci). Lysate was mixed by vortexing and sonicated for 2 minutes at 100 amplitude with QSonica Q700 sonicator (Newton, CT). Lysates were then centrifuged at 14,000 rpm for 15 min at 4°C. Supernatant was transferred to a fresh vial and total protein contents were estimated by DC Protein Assay kit (BioRad) and equal amounts of proteinswere resolved on 4-15% gradient (SDS)-polyacrylamide gel (Bio-Rad). The proteins were then electro-transferred for 10 minutes onto polyvinylidene-difluoride (PVDF) membrane using Transblot Turbo device (Bio-Rad). After blocking with 5% non-fat powdered milk in Tris-buffered saline (TBS, 0.1 M, pH = 7.4), blots were incubated with primary antibody at 4°C overnight. The following primary antibodies were used: p300 (NM11) from Santa Cruz Biotechnology (Santa Cruz, CA); Acetyl-Lysine from Abcam (Cambridge, United Kingdom) and CBP, H3K9, H3K18, H3K27, total Histone3 Cleaved Caspase-3 and β-actin from Cell Signaling Technology (Danvers, MA). After each step, blots were washed three times with Tween (0.1%)-Tris-buffer saline (TBS-T). Goat anti-mouse and anti-rabbit secondary antibody conjugated to horseradish peroxidase (GE Healthcare, Chicago, Illinois) were used, and signal was generated with the ECL2 western blotting substrate (Pierce Biotechnology, Rockford, IL) on HyBlot CL autoradiographic film (Thomas Scientific, Swedesboro, NJ). Protein abundance of β-actin served as a control for protein loading in each lane.

**Immunoprecipitation**

Cells were grown in 15cm dishes until 75% confluency and lysed using 1ml Pierce IP lysis buffer (Thermo) containing protease and phosphatase inhibitors (Lysates were sonicated for 2 minutes at 100% amplitude and cell debris was removed by centrifugation at 14K rpm for 15 minutes. Protein quantification was estimated using DC protein assay kit. 500 micrograms of each sample were immunoprecipitated with 5ul CBP antibody (Cell Signaling Tech), 25ul BRCA1 antibody (Santa Cruz) or anti-acetyl lysine affinity beads (Cytoskeleton Inc.) and incubated overnight rotating at 4C. For CBP and BRCA1 IP, 50ul of 100mg/ml Protein A Sepharose beads (GE Healthcare) were added and rotated at 4C for 2h the following day. Three 1ml washes with IP lysis buffer were used to isolate the precipitate and samples were boiled in 25ul 2X SDS loading buffer for 7 minutes and loaded into 4-15% polyacrylamide gels (BioRad).

1.	Carugo, A. *et al.* In Vivo Functional Platform Targeting Patient-Derived Xenografts Identifies WDR5-Myc Association as a Critical Determinant of Pancreatic Cancer. *Cell Rep.* **16**, 133–147 (2016).

2.	Birmingham, A. *et al.* Statistical Methods for Analysis of High-Throughput RNA Interference Screens. *Nat. Methods* **6**, 569–575 (2009).
