## Supplementary Tables and Figures for "Inhibition of histone acetyltranserase function radiosensitizes CREBBP/EP300 mutants via repression of homologous recombination, potentially targeting a novel gain of function"

| **Cell Line** | **CREBBP status** | **EP300 status** | **P53 status** | **HPV status** |
| --- | --- | --- | --- | --- |
| Cal27 | T2390N | WT | H193L | - |
| UM-SCC-22a | Q1773X | WT | Y220C | - |
| UM-SCC-25 | WT | P1780fs | Splice site | - |
| HN5 | Q2158P | R1055X | C238S | - |
| UM-SCC-47 | Q1092X | WT | WT | + |
| HN31 | WT | WT | C176F, A161S | - |
| HN30 | WT | Q793splice, | WT | - |
|  |  | V1594splice |  |  |
| UPCI:SCC152 | WT | WT | WT | + |
| FaDu | WT | WT | R248L | - |
| UM-SCC-1 | WT | A4241G | Splice site | - |
| H520 | R1446C | WT | W146X |  |
| H2228 | WT | A2251T | Q331X |  |
| Calu6 | WT | WT | R196X |  |
| HEK-293T | WT | WT | WT |  |

**Supplementary Table 1. Characteristics of the cell lines used in this study.**

| **Targetable** |  |  |  | **DNA Damage** |  |  |  |  |  |  |  |
| --- | --- | --- | --- | --- | --- | --- | --- | --- | --- | --- | --- |
| ABL1 | FGFR4 | MAPK7 | PRKDC | ABCF2 | CDK6 | EXO1 | MDM4 | POLD3 | REV3L | TCEA1 | XRCC4 |
| ABL2 | FGR | MAPK8 | PSMA1 | ABL1 | CDK7 | FANCA | MGMT | POLDIP2 | REXO2 | TDG | XRCC5 |
| AKT1 | FLT1 | MAPK9 | PSMB1 | AIFM1 | CDKN1A | FANCB | MLH1 | POLE | RFC1 | TDP1 | XRCC6 |
| AKT2 | FLT3 | MAPT | PSMD1 | ALKBH1 | CDKN2A | FANCC | MLH3 | POLE2 | RFC2 | TERF1 | BP1 |
| AKT3 | FLT4 | MCL1 | PTCH1 | ANKRD17 | CDKN2B | FANCD2 | MMS19 | POLG | RFC3 | TERF2 | XRN2 |
| ALK | FRK | MDM2 | PTGS2 | ANTXR1 | CDKN2D | FANCE | MNAT1 | POLG2 | RFC4 | TERT | YBX1 |
| AR | FYN | MET | PTK2 | APAF1 | CDT1 | FANCF | MPG | POLH | RFC5 | TFF2 | ZAK |
| ATM | GLS | MKNK1 | PTPN11 | APEX1 | CEBPG | FANCG | MRE11A | POLI | RFWD2 | TGFB1 | ZW10 |
| ATR | GSK3A | MS4A1 | PTPN6 | APEX2 | CENPF | FANCI | MSH2 | POLK | RINT1 | TIMELESS | ZWINT |
| AURKA | GSK3B | MTOR | RAC1 | APTX | CETN2 | FANCL | MSH3 | POLL | RNF168 | TIPIN |  |
| AURKB | HDAC1 | MYC | RAF1 | ASF1A | CHAF1A | FANCM | MSH4 | POLM | RNF8 | TNFRSF10B |  |
| AURKC | HDAC2 | NAMPT | RARA | ATM | CHAF1B | FEN1 | MSH5 | POLN | RPA1 | TNP1 |  |
| AXL | HDAC3 | NFKB1 | RARB | ATR | CHEK1 | FOXN3 | MSH6 | POLQ | RPA2 | TOP1 |  |
| BCL2 | HDAC6 | NOTCH1 | RARG | ATRIP | CHEK2 | FUS | MTOR | POLR2G | RPA3 | TOP2A |  |
| BCR | HDAC8 | NR2C2 | RET | ATRX | CHFR | GADD45A | MUS81 | POT1 | RPA4 | TOPBP1 |  |
| BIRC5 | HSP90AA1 | NTRK1 | ROCK1 | ATXN3 | CIB1 | GADD45B | MUTYH | PPM1D | RPAIN | TP53 |  |
| BLK | IDH1 | NUDT1 | ROCK2 | BAI1 | CIDEA | GADD45G | NAE1 | PRIM1 | RPL13A | TP53BP1 |  |
| BMX | IDH2 | P4HB | RPL30 | BARD1 | CRY1 | GTF2E2 | NBN | PRIM2 | RPL30 | TP73 |  |
| BRAF | IGF1R | PAK1 | RPS6KB1 | BCL2 | CRY2 | GTF2H1 | NCOA6 | PRKCG | RPRM | TREX1 |  |
| BRD4 | IKBKE | PAK4 | RRM1 | BIRC5 | CSNK1D | GTF2H2 | NEIL1 | PRKDC | RPS27A | TREX2 |  |
| BTK | IL1B | PARP1 | RXRA | BLM | CSNK1E | GTF2H2B | NEIL2 | PSMA1 | RRM1 | TRIAP1 |  |
| CASP3 | IL6 | PARP2 | RXRB | BRCA1 | CUL4A | GTF2H3 | NEK11 | PSME4 | RRM2 | TRRAP |  |
| CCR5 | IL6R | PARP3 | SGK3 | BRCA2 | CUL4B | GTF2H4 | NHEJ1 | PTEN | RRM2B | TTK |  |
| CD274 | INSR | PDGFRA | SMO | BRIP1 | CYCS | GTF2H5 | NME2 | PTTG1 | RUVBL1 | TXN |  |
| CD52 | IRAK4 | PDGFRB | SRC | BRSK1 | DCLRE1A | GTSE1 | NTHL1 | RAD1 | RUVBL2 | UBA52 |  |
| CDK1 | ITK | PDK1 | STAT3 | BTG2 | DCLRE1B | H2AFX | NUDT1 | RAD17 | SEMA4A | UBB |  |
| CDK2 | JAK1 | PGD | SYK | BUB1 | DCLRE1C | HDAC4 | OGG1 | RAD18 | SESN1 | UBE2A |  |
| CDK4 | JAK2 | PIGF | TBK1 | BUB1B | DDB1 | HMGB2 | OXR1 | RAD21 | SETX | UBE2B |  |
| CDK6 | JAK3 | PIK3CA | TEC | CCNA2 | DDB2 | HPRT1 | PALB2 | RAD23A | SF3B3 | UBE2I |  |
| CDK7 | KDM1A | PIK3CB | TEK | CCNB1 | DDIT3 | HUS1 | PARG | RAD23B | SHFM1 | UBE2N |  |
| CDK9 | KDR | PIK3CD | TNF | CCNB2 | DDX11 | IGHMBP2 | PARP1 | RAD50 | SHISA5 | UBE2T |  |
| CHEK1 | KIT | PIK3CG | TNFRSF8 | CCNB3 | DKC1 | ING1 | PARP2 | RAD51 | SIAH1 | UBE2V1 |  |
| CHEK2 | LAP3 | PIM1 | TNFSF11 | CCND1 | DLGAP5 | INPPL1 | PARP3 | RAD51B | SIRT1 | UBE2V2 |  |
| CREBBP | LCK | PIM2 | TNFSF13B | CCND2 | DMC1 | IP6K3 | PARP4 | RAD51C | SLK | UNG |  |
| CTNNB1 | LDHA | PIM3 | TOP1 | CCND3 | DNA2 | KAT2A | PCNA | RAD51D | SLX4 | UPF1 |  |
| DHFR | LUC | PLK1 | TOP2A | CCNE1 | DNTT | KAT5 | PERP | RAD52 | SMC1A | USP1 |  |
| DOT1L | LYN | PORCN | TOP2B | CCNE2 | DUT | KNTC1 | PML | RAD54B | SMC2 | UVRAG |  |
| DRD2 | MAP2 | PPM1D | TRIM24 | CCNG1 | E2F1 | LIG1 | PMS1 | RAD54L | SMC3 | VCP |  |
| EGFR | MAP2K1 | PRKAA1 | TRPV1 | CCNG2 | EI24 | LIG3 | PMS2 | RAD9A | SMC6 | WDR33 |  |
| EHMT2 | MAP2K2 | PRKCA | TUBB | CCNH | EME1 | LIG4 | PMS2L2 | RB1 | SOD1 | WRAP53 |  |
| EIF4E | MAP3K14 | PRKCB | TXN | CCNO | EP300 | LRIG1 | PMS2P1 | RBBP4 | SPO11 | WRN |  |
| EPHA2 | MAP3K8 | PRKCD | TYMS | CDC25A | ERCC1 | LUC | PMS2P3 | RBBP8 | SSBP1 | WRNIP1 |  |
| ERBB2 | MAP4 | PRKCE | TYRO3 | CDC25B | ERCC2 | MAD2L1 | PMS2P4 | RBM14 | STEAP3 | XAB2 |  |
| ESR1 | MAPK1 | PRKCG | VEGFA | CDC25C | ERCC3 | MAD2L2 | PMS2P5 | RECQL | SUMO1 | XPA |  |
| ESR2 | MAPK11 | PRKCH | WEE1 | CDC6 | ERCC4 | MBD4 | PNKP | RECQL4 | SUPT3H | XPC |  |
| EZH2 | MAPK12 | PRKCI | WHSC1 | CDK1 | ERCC5 | MCM8 | POLA1 | RECQL5 | TADA3 | XRCC1 |  |
| FGFR1 | MAPK13 | PRKCQ | XIAP | CDK2 | ERCC6 | MDC1 | POLB | RELA | TAF2 | XRCC2 |  |
| FGFR2 | MAPK14 | PRKCSH | XPO1 | CDK4 | ERCC8 | MDM2 | POLD1 | REV1 | TAF5L | XRCC3 |  |
| FGFR3 | MAPK3 | PRKCZ |  |  |  |  |  |  |  |  |  |

**Supplementary Table 2. Targets of the in vivo shRNA screening.**

| **Gene** | **Average untreated** | **Average XRT** | **Gene** | **Average untreated** | **Average XRT** | **Gene** | **Average untreated** | **Average XRT** |
| --- | --- | --- | --- | --- | --- | --- | --- | --- |
| ABL1 | -0.549 | -0.858 | KDM1A | -0.666 | -1.080 | RUVBL1 | -1.964 | -2.683 |
| AKT1 | -0.720 | -1.222 | LCK | -0.917 | -1.401 | SEMA4A | -0.626 | -0.947 |
| ANTXR1 | -0.909 | -1.230 | MAD2L1 | -1.303 | -1.534 | SF3B3 | -1.522 | -1.704 |
| ATM | -1.122 | -1.278 | MAP3K8 | -1.906 | -2.190 | SMC1A | -2.111 | -2.918 |
| ATRIP | -1.806 | -2.370 | MAPK12 | -0.653 | -1.095 | SMC6 | -2.018 | -2.506 |
| ATXN3 | -0.700 | -0.945 | MCL1 | -1.321 | -1.708 | SPO11 | -0.683 | -1.254 |
| AURKB | -0.900 | -1.042 | MDM2 | -0.691 | -1.165 | SSBP1 | -0.946 | -1.442 |
| BCL2 | -0.629 | -1.051 | MKNK1 | -0.553 | -0.901 | SYK | -0.742 | -0.924 |
| BLK | -0.864 | -1.115 | MSH5 | -0.513 | -1.032 | TOP2A | -0.660 | -0.976 |
| BLM | -0.607 | -0.896 | MTOR | -0.774 | -0.949 | TOPBP1 | -0.816 | -1.007 |
| BRAF | -1.084 | -1.349 | MTOR | -0.934 | -1.337 | TP53BP1 | -0.674 | -1.477 |
| BRCA2 | -0.600 | -1.059 | NEIL2 | -1.800 | -2.093 | TTK | -0.515 | -0.954 |
| BRD4 | -1.228 | -1.553 | NOTCH1 | -0.980 | -1.167 | TXN | -0.747 | -1.029 |
| BTG2 | -0.964 | -1.090 | NUDT1 | -1.261 | -1.598 | TYMS | -0.743 | -0.934 |
| BTK | -0.636 | -0.945 | OXR1 | -0.756 | -0.867 | UBB | -0.750 | -0.900 |
| CCNB1 | -1.136 | -1.300 | PAK4 | -0.916 | -1.113 | UBE2A | -0.670 | -0.911 |
| CCNO | -1.803 | -2.782 | PARG | -1.568 | -1.961 | UBE2V2 | -2.217 | -2.590 |
| CDC25A | -1.057 | -1.372 | PARP4 | -0.679 | -1.002 | VCP | -1.497 | -1.647 |
| CDC25B | -0.713 | -0.867 | PERP | -0.296 | -0.814 | XIAP | -1.729 | -2.352 |
| CDK2 | -0.843 | -1.069 | PIGF | -0.769 | -0.894 | XPA | -0.645 | -0.863 |
| CDK4 | -0.958 | -1.315 | PIK3CA | -2.798 | -3.597 | XRCC5 | -1.064 | -1.238 |
| CENPF | -0.781 | -1.071 | PMS1 | -0.948 | -1.179 | XRCC6 | -1.431 | -1.732 |
| CHEK1 | -1.352 | -1.741 | PMS2P3 | -0.357 | -0.904 | YBX1 | -1.912 | -2.264 |
| CIB1 | -0.803 | -0.942 | PMS2P5 | -0.765 | -1.282 | ZW10 | -0.977 | -1.289 |
| CREBBP | -0.588 | -0.899 | POLA1 | -1.097 | -1.618 | ZWINT | -0.963 | -1.829 |
| CUL4A | -0.670 | -1.144 | POLG | -1.016 | -1.311 |  |  |  |
| DLGAP5 | -0.810 | -0.971 | POLL | -1.115 | -1.416 |  |  |  |
| DRD2 | -0.608 | -1.037 | PRKCQ | -0.619 | -0.857 |  |  |  |
| DUT | -1.763 | -2.038 | PRKDC | -0.942 | -1.266 |  |  |  |
| EHMT2 | -0.688 | -1.015 | PRKDC | -1.508 | -1.807 |  |  |  |
| EP300 | -0.589 | -1.010 | PTK2 | -1.545 | -1.857 |  |  |  |
| EPHA2 | -1.264 | -1.537 | RAD1 | -0.805 | -1.076 |  |  |  |
| ERCC6 | -1.241 | -1.389 | RAD51 | -1.151 | -1.443 |  |  |  |
| ERCC8 | -0.962 | -1.179 | RAD51C | -0.797 | -0.960 |  |  |  |
| ESR1 | -0.817 | -1.396 | REXO2 | -0.907 | -1.118 |  |  |  |
| FANCD2 | -0.727 | -1.157 | RFC2 | -0.881 | -1.000 |  |  |  |
| FANCE | -0.840 | -1.435 | RNF168 | -0.866 | -1.118 |  |  |  |
| FANCL | -0.973 | -1.144 | RPA1 | -1.149 | -1.277 |  |  |  |
| FGFR4 | -1.163 | -1.410 | RPA2 | -2.374 | -2.761 |  |  |  |
| GTF2E2 | -1.113 | -1.232 | RPRM | -0.587 | -0.989 |  |  |  |
| JAK2 | -0.693 | -1.071 | RPS6KB1 | -0.663 | -0.969 |  |  |  |
| KAT2A | -1.173 | -1.353 | RRM2B | -0.827 | -0.927 |  |  |  |

**Supplementary Table 3. Targets from XRT vs. untreated tumor analysis.**

| **Site** | **N** | **%** |
| --- | --- | --- |
| Oral Cavity | 152 | 55.1% |
| Oropharynx | 56 | 20.3% |
| Larynx/hypopharynx | 68 | 24.6% |
| **Nodal stage** |  |  |
| 0 | 101 | 36.9% |
| 1 | 53 | 19.3% |
| 2x | 3 | 1.1% |
| 2a | 14 | 5.1% |
| 2b | 61 | 22.3% |
| 2c | 29 | 10.6% |
| 3 | 5 | 1.8% |
| Unknown | 8 | 2.9% |
| **Tumor stage** |  |  |
| 1 | 15 | 5.5% |
| 2 | 60 | 21.9% |
| 3 | 78 | 28.5% |
| 4 | 115 | 42.0% |
| Unknown | 6 | 2.2% |

**Supplementary Table 4. Clinical characteristics of the patients in the Head and Neck TCGA cohort treated with radiation.**

| **Site** | **N** | **%** |
| --- | --- | --- |
| Oral Cavity | 60 | 63.8% |
| Oropharynx | 4 | 4.3% |
| Larynx/hypopharynx | 30 | 31.9% |
| **Nodal stage** |  |  |
| 1 | 44 | 46.8% |
| 2x | 20 | 21.3% |
| 2a | 1 | 1.1% |
| 2b | 2 | 2.1% |
| 2c | 13 | 13.8% |
| 3 | 8 | 8.5% |
| Unknown | 2 | 2.1% |
|  | 4 | 4.3% |
| **Tumor stage** |  |  |
| 1 | 1 | 1.1% |
| 2 | 12 | 12.8% |
| 3 | 23 | 24.5% |
| 4 | 56 | 59.6% |
| Unknown | 2 | 2.1% |

**Supplementary Table 5. Clinical characteristics of patients in the subset analysis.**

|  | **Lung SCC** | |  |  | **Cervix SCC** | |
| --- | --- | --- | --- | --- | --- | --- |
| **Overall stage** | **N** | **%** |  | **Overall stage** | **N** | **%** |
| IA | 2 | 3.3 |  | IB1 | 10 | 15.2 |
| IB | 4 | 6.6 |  | IB2 | 6 | 9.1 |
| II | 1 | 1.6 |  | IIA | 2 | 3.0 |
| IIA | 6 | 9.8 |  | IIA2 | 1 | 1.5 |
| IIB | 14 | 23.0 |  | IIB | 22 | 33.3 |
| III | 1 | 1.6 |  | III | 1 | 1.5 |
| IIIA | 26 | 42.6 |  | IIIA | 3 | 4.5 |
| IIIB | 7 | 11.5 |  | IIIB | 11 | 16.7 |
|  |  |  |  | IVA | 3 | 4.5 |
| **Nodal stage** |  |  |  | IVB | 7 | 10.6 |
| N0 | 16 | 26.2 |  |  |  |  |
| N1 | 17 | 27.9 |  | **Nodal stage** |  |  |
| N2 | 22 | 36.1 |  | N0 | 11 | 16.7 |
| N3 | 5 | 8.2 |  | N1 | 8 | 12.1 |
| NX | 1 | 1.6 |  | NX | 47 | 71.2 |
| **Tumor stage** |  |  |  | **Tumor stage** |  |  |
| T1 | 4 | 6.6 |  | T1b | 2 | 3.0 |
| T2 | 24 | 39.3 |  | T1b1 | 6 | 9.1 |
| T2a | 6 | 9.8 |  | T1b2 | 4 | 6.1 |
| T2b | 4 | 6.6 |  | T2 | 2 | 3.0 |
| T3 | 18 | 29.5 |  | T2a | 2 | 3.0 |
| T4 | 5 | 8.2 |  | T2a2 | 2 | 3.0 |
|  |  |  |  | T2b | 18 | 27.3 |
|  |  |  |  | T3 | 2 | 3.0 |
|  |  |  |  | T3a | 2 | 3.0 |
|  |  |  |  | T3b | 10 | 15.2 |
|  |  |  |  | T4 | 5 | 7.6 |
|  |  |  |  | Tis | 1 | 1.5 |
|  |  |  |  | TX | 10 | 15.2 |
|  |  |  |  | **Grade** |  |  |
|  |  |  |  | G1 | 3 | 4.5 |
|  |  |  |  | G2 | 28 | 42.4 |
|  |  |  |  | G3 | 23 | 34.8 |
|  |  |  |  | GX | 12 | 18.2 |

**Supplementary Table 6. Clinical characteristics of patients in the lung and cervix SCC cohorts.**

**Supplemental figures.**

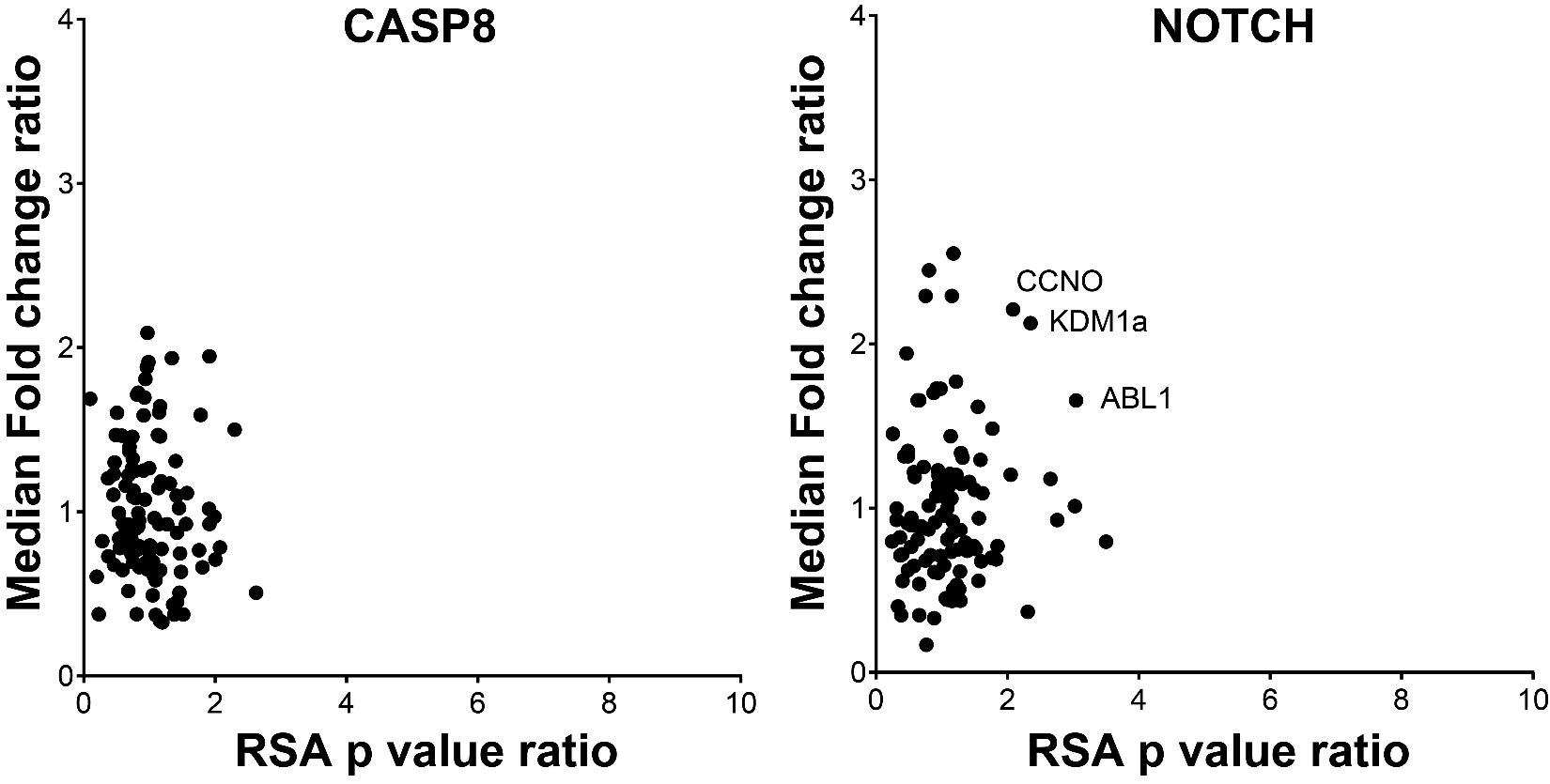

**Supplementary Figure 1.** Ratio of CASP8 or NOTCH mutant vs. wild type for target fold change (y-axis) and RSA log p-value (x-axis) for radiosensitizing targets selected from Fig. 1.

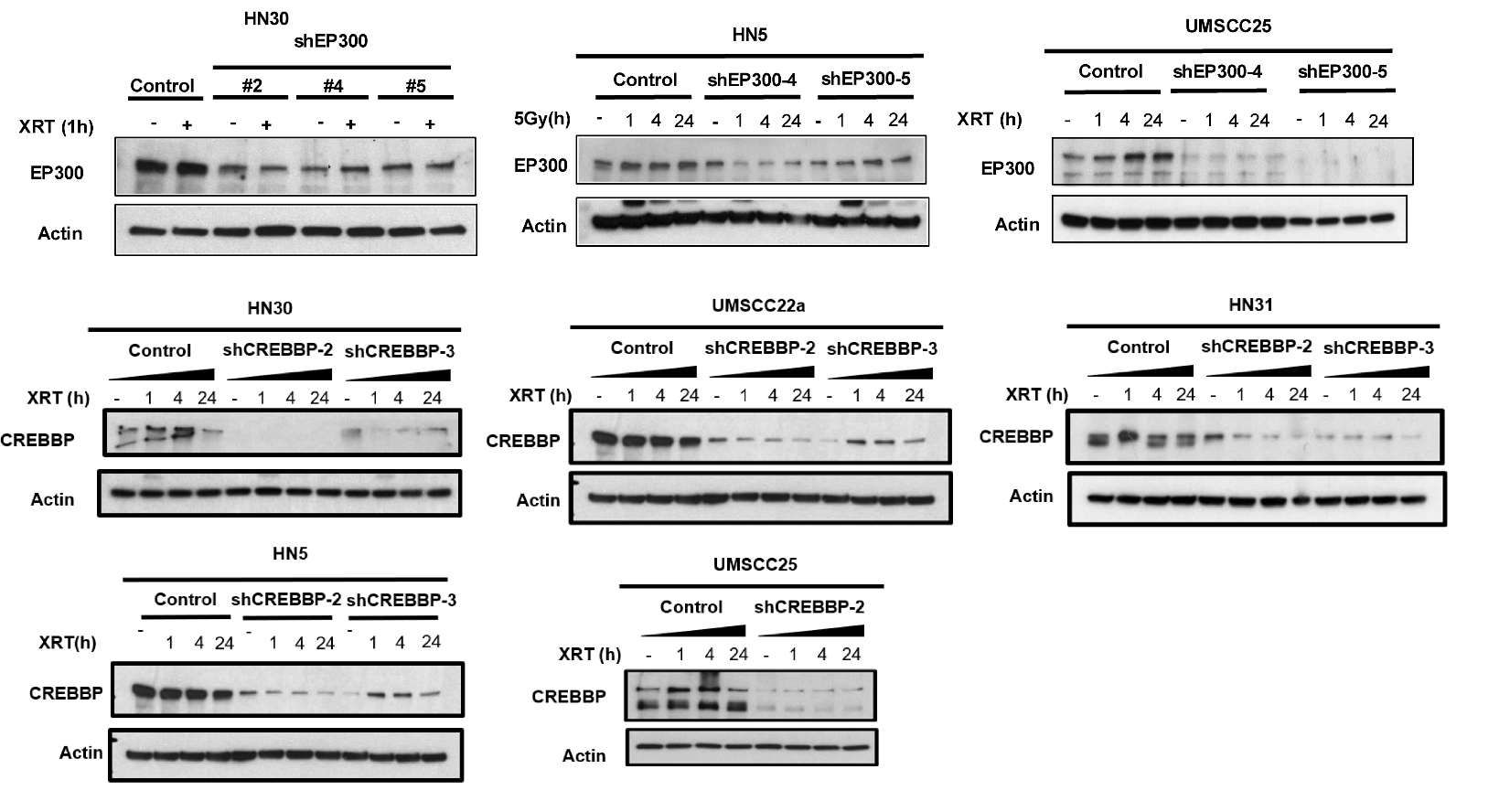

**Supplementary Figure 2.** Immunoblot for CREBBP and EP300 in shRNA knockdown cells.

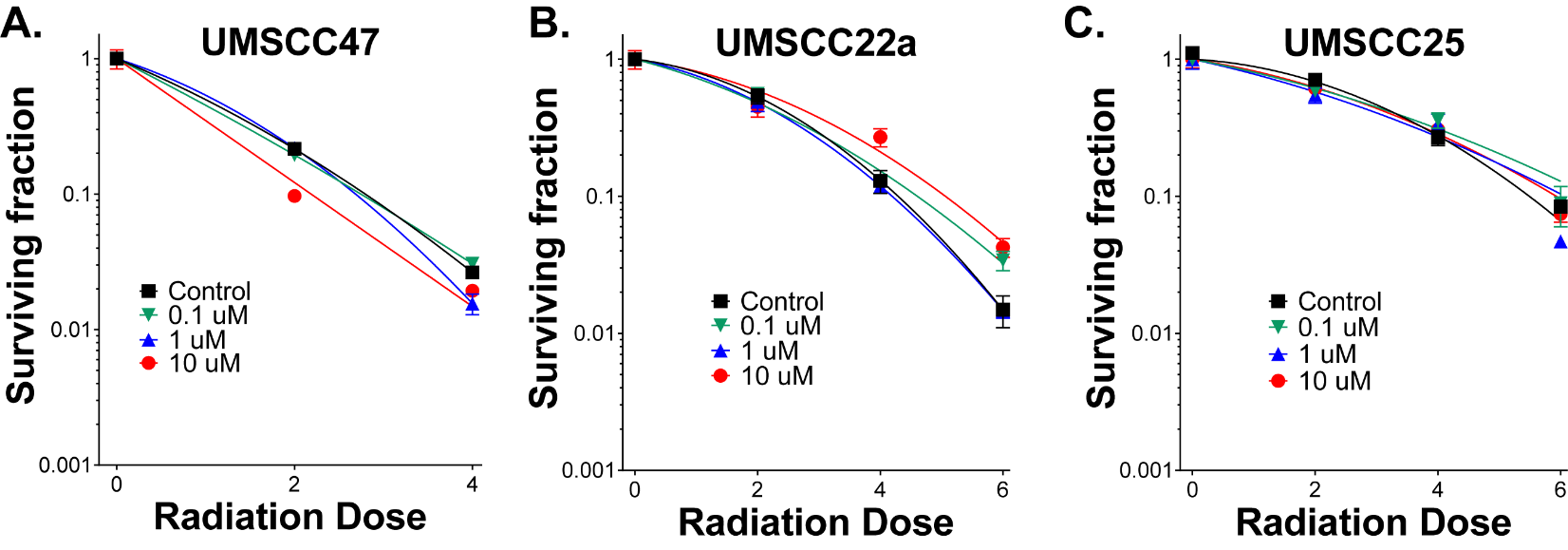

**Supplementary Figure 3.** A-C) Clonogenic survival data following treatment with GNE-272, a bromodomain specific inhibitor for both CREBBP and EP300.

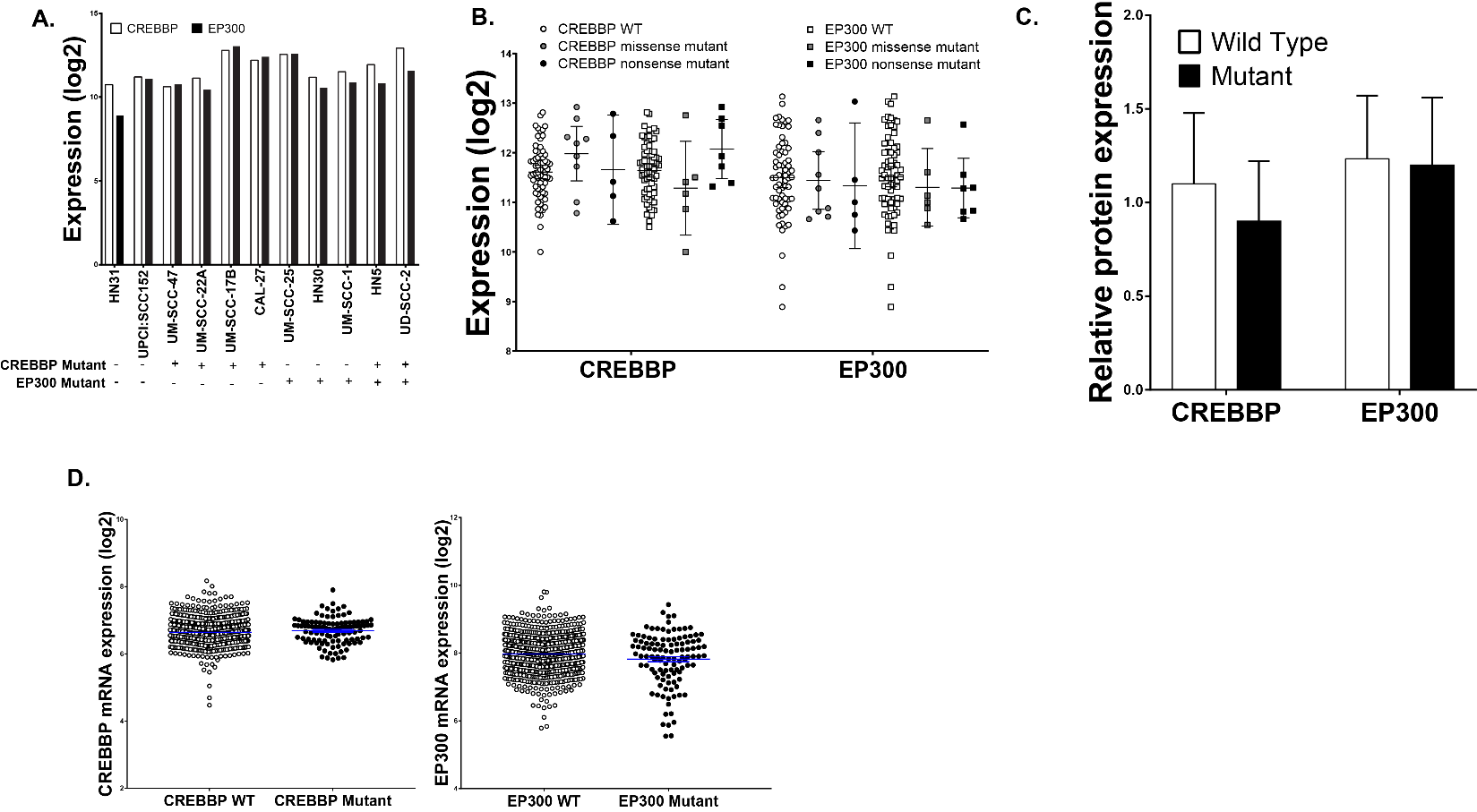

**Supplementary Figure 4.** A & B) CREBBP and EP300 mRNA expression in the cell lines used in this study (A) and in a total of 82 HNSCC cell lines described previously (B). C) densitometry for CREBBP and EP300 from the cell lines used in this study. D) CREBBP and EP300 mRNA expression all available tumors from the Head and Neck TCGA.
